# A tunable receptor separates root barrier formation from nutrient signaling through ligand-perception states

**DOI:** 10.64898/2026.07.06.736668

**Authors:** Yuanyuan Zhang, Sebastian Samwald, Anika Schröder, Sara Stolze, Swati Mahiwal, Tianquan Lu, Jakub Rzemieniewski, Martin Stegmann, Hirofumi Nakagami, Defeng Shen, Tonni Grube Andersen

## Abstract

In roots, the endodermis controls nutrient entry by forming a barrier known as the Casparian strip^1–3^. Yet how barrier-associated processes interface with systemic signaling remains unclear. Here, we show that the receptor kinase SCHENGEN3 links local barrier surveillance to systemic nutrient signaling, with outputs that depend on the effective state of ligand perception. Unlike in *Arabidopsis thaliana*, SCHENGEN3 activation in *Lotus japonicus* requires a distinct cellular competence state and cannot be triggered by exogenous ligands alone, revealing evolutionary divergence in pathway deployment. Cross-species complementation uncouples systemic nitrogen signaling from Casparian strip formation, while transcriptomic and phosphoproteomic analyses reveal largely non-overlapping signaling- and barrier-associated programs that differ between agar and agricultural soil conditions. Mechanistically, receptor-ligand comparisons, engineered receptor variants, and co-receptor mutant analyses show that systemic nitrogen signaling is retained in receptor-perception states that are insufficient to support full Casparian strip establishment. Together, these findings reveal how a shared receptor module can separate developmental and physiological outputs by linking receptor perception state to output specificity.

## Main text

Plants continuously adjust nutrient uptake and distribution in response to changing environmental conditions. In roots, this process is shaped by the endodermis — a specialized cell layer that surrounds the vasculature and controls apoplastic access to the inner tissues (the stele)^1–3^. This control is mainly mediated by the Casparian strip (CS), which blocks apoplastic flow between the endodermal cells, and thereby forces solutes to enter the stele through plasma membrane-localized active transport systems^1–3^. Barrier function is further reinforced by suberin deposition at the endodermal cell surface, which limits exchange with the surrounding apoplast^1,3,4^. Although this structural filtration role is well established, we know little about how endodermal barrier-associated pathways are integrated with systemic nutrient signaling. In the model plant *Arabidopsis thaliana* (hereafter *Arabidopsis*), the SCHENGEN (SGN) pathway monitors CS integrity and, upon disruption, promotes efficient re-sealing through perception of CASPARIAN STRIP INTEGRITY FACTOR (CIF) peptides by the leucine-rich receptor-like kinase SGN3 (Refs.^5,6^). Loss of CS integrity activates SGN signaling, resulting in compensatory lignification and enhanced suberization^4,7,8^. CIF peptides are required for SGN pathway activation in *Arabidopsis* and rice (*Oryza* ssp.)^9^, yet how far this ligand-recognition logic and downstream signaling architecture are conserved remains unknown. In addition to its role in root barrier surveillance, components of the SGN pathway are implicated in nutrient-dependent shoot growth responses, suggesting that SGN signaling may also contribute to systemic physiology^10^. Whether these functions are mechanistically linked and how a single receptor coordinates structurally distinct outputs remain unresolved.

Here, we show that SGN3 links ligand perception to distinct physiological outputs through output-specific receptor-perception states. Using comparative genetics in *Lotus japonicus* (hereafter *Lotus*) and *Arabidopsis*, we find that SGN pathway activation in *Lotus* requires a MYB36-dependent competence state involving expression of core SGN pathway components, revealing divergence in pathway deployment. Cross-species complementation uncouples SGN3-dependent CS formation from systemic nitrogen signaling, while engineered receptor variants demonstrate that these outputs depend on distinct receptor-perception states. Transcriptomic and phosphoproteomic analyses further show that SGN3-dependent nitrogen signaling is most prominent under agricultural soil conditions and comparatively limited in standard agar assays. This nitrogen branch engages the canonical C-TERMINALLY ENCODED PEPTIDE (CEP)–CEP RECEPTOR (CEPR) signaling module, linking local endodermal receptor activity to systemic nutrient responses. Together, this establishes SGN3 as a tunable receptor that links ligand perception to output selection, thereby separating developmental barrier formation from systemic nutrient signaling.

### SGN activation in *Lotus* requires a MYB36-dependent competence state

In *Arabidopsis*, disruption of CS formation activates the SGN pathway and triggers compensatory lignification and early endodermal suberization^4,7,8^. In *Lotus*, however, these hallmark SGN responses were absent in two independent *Ljmyb36* KO alleles, despite severe CS defects (Extended Data Fig. 1a,b)^11^. This suggested that SGN pathway activation is regulated differently in *Lotus*, prompting us to test whether differences in ligand availability or composition could explain the lack of response. We identified two putative CIF peptide-encoding genes in the *Lotus* genome^12^. One candidate showed strong root-enriched expression, approximately 20-fold higher than the second candidate, and was therefore designated *LjCIF2*, whereas the lower-expressed candidate was designated *LjCIF1* according to *Arabidopsis* nomenclature and expression patterns (Extended Data Fig. 1c,d). Sequence analysis revealed that both predicted mature *Lotus* CIF peptides contain two tyrosine residues with potential sulfation sites, whereas *Arabidopsis* CIF peptides carry only one conserved sulfated tyrosine (Extended Data Fig. 1c). We therefore tested whether the sulfation state of LjCIF2 affects SGN pathway activation using single (LjCIF2_sTyr2_) and double (LjCIF2_sTyr2Tyr5_) sulfated LjCIF2 variants.

When applied to *Arabidopsis* roots, both LjCIF2 variants induced SGN-associated responses, including ectopic lignification and increased suberization (Extended Data Fig. 1e,g). However, the double-sulfated variant was less active than the single-sulfated form and weaker than native AtCIF2_sTyr2_ (Extended Data Fig. 1e), indicating that sulfation state modulates ligand potency. In contrast, none of the tested CIF peptides, including AtCIF2_sTyr2_, LjCIF2_sTyr2_, and LjCIF2_sTyr2Tyr5_ induced suberization or lignification in *Lotus* roots, even at five-fold higher concentrations (Extended Data Fig. 1e,g). This lack of response was not explained by impaired peptide delivery, as hydroponic peptide application gave similar results and fluorescently labeled AtCIF2 was readily detected in the endodermis of differentiated *Lotus* roots (Extended Data Fig. 1f-h). Thus, exogenous CIF peptide application is not sufficient to activate SGN-associated cell-wall deposition outputs in *Lotus* under the tested conditions (Fig.1a).

We next examined whether CS-defective *Lotus* roots activate transcriptional SGN responses. In *Arabidopsis*, CS defects induce SGN-dependent compensatory programs, whereas combined loss of MYB36 and SGN3 suppresses this activation^13^. To test the corresponding relationship in *Lotus*, we generated a *Ljmyb36-2* × *Ljsgn3-2* double mutant and compared its root transcriptome with wild type (WT) plants and the respective single mutants. Consistent with the lack of SGN-associated cell-wall responses, canonical genes associated with SGN pathway activation, including those linked to immune responses and phenylpropanoid biosynthesis, were not induced in *Ljmyb36-2* roots (Fig. 1b). *Lotus* orthologs of *Arabidopsis* genes associated with CS formation, including *LjCASPARIAN STRIP MEMBRANE DOMAIN PROTEINs* (*LjCASPs*)^14^, *LjENHANCED SUBERIN 1* (*LjESB1*)^7^, and *LjPEROXIDASE64* (*LjPER64*)^15^, were downregulated in CS-defective *Lotus* mutants compared with WT plants (Fig. 1c and Extended Data Fig. 2a). Consistent with reduced SGN activity, the *Ljmyb36-2* × *Ljsgn3-2* double mutant clustered closer to *Ljmyb36-*2 than to *Ljsgn3-2* or WT in principal component space, indicating that loss of *LjSGN3* has only a minor transcriptional impact in the absence of *MYB36* (Extended Data Fig. 2b). This was further supported by the reduced expression of core SGN components. *LjSGN1* and *LjSGN3* were downregulated in *Ljmyb36-2* roots, and promoter-reporter lines showed little or no detectable activity in the CS-establishment zone (Extended Data Fig. 2c,d). Moreover, endodermal expression of *LjSGN1* and *LjSGN3* under the *LjSCR* promoter partially restored early suberization in the *Ljmyb36-2* background (Extended Data Fig. 2e-g). Together, these results show that *Lotus* CIF peptides can act as SGN3 ligands, but ligand availability alone is insufficient to trigger canonical SGN outputs in *Lotus*. Instead, SGN activation in *Lotus* requires a MYB36-dependent competence state that includes expression of core SGN signaling components (Fig. 1d).

**Fig. 1.**
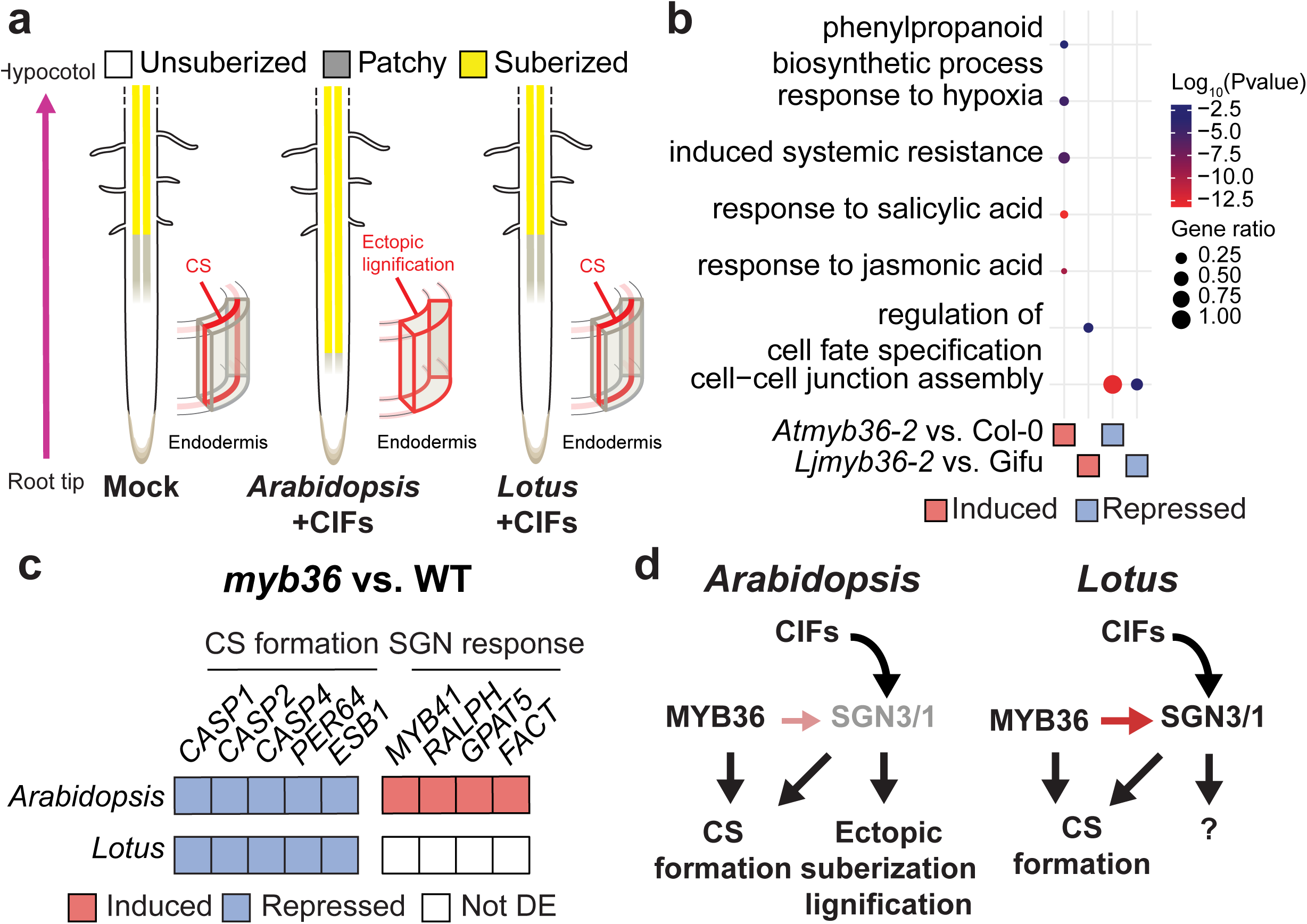
Activation of the *Lotus* SGN pathway requires MYB36. **a**, Schematic illustration of distinct SGN responses triggered by exogenous CIF peptides in *Arabidopsis* and *Lotus*. **b**, Dotplot showing Gene Ontology terms enriched among induced and repressed differentially expressed genes (FDR-adjusted *P* value < 0.05, absolute fold change > 2) in *Atmyb36-2* and *Ljmyb36-2* roots relative to their respective wild type (WT) controls. **c**, Heatmap showing the expression of genes associated with Casparian strip (CS) formation and SGN response in *Arabidopsis* and *Lotus myb36* mutant roots relative to their WT. Red: induced (FDR-adjusted *P* value < 0.05, fold change >2); blue: repressed (FDR-adjusted *P* value < 0.05, fold change < -2); white: not differentially expressed (DE). **d**, Schematic illustration showing activation of the SGN pathway requires MYB36 in *Lotus*, yet unknown downstream outputs by exogenous CIF peptides.

### Cologne agricultural soil reveals a separable SGN3-dependent nitrogen-signaling branch

We next asked whether SGN1 and SGN3 functions are conserved between *Arabidopsis* and *Lotus* despite their divergent SGN activation logic. Consistent with conserved roles in barrier formation, both AtSGN1 and LjSGN1 restored CS function in *Atsgn1-1* (Fig. 2a). In contrast, only AtSGN3 complemented CS formation in *Atsgn3-3*, whereas *LjSGN3*-expressing lines retained a defective barrier (Fig. 2a). We recently showed that the *Atsgn3-3* mutant displays decreased nitrogen-induced growth and anthocyanin accumulation under Cologne agricultural soil (CAS) conditions^10^. Strikingly, both phenotypes were restored by AtSGN3 and LjSGN3, although LjSGN3 failed to rescue the defective CS phenotype of *Atsgn3-3* under CAS conditions (Fig. 2b,c and Extended Data Fig. 3). Thus, SGN3-dependent nitrogen responses can be genetically uncoupled from CS integrity.

**Fig. 2.**
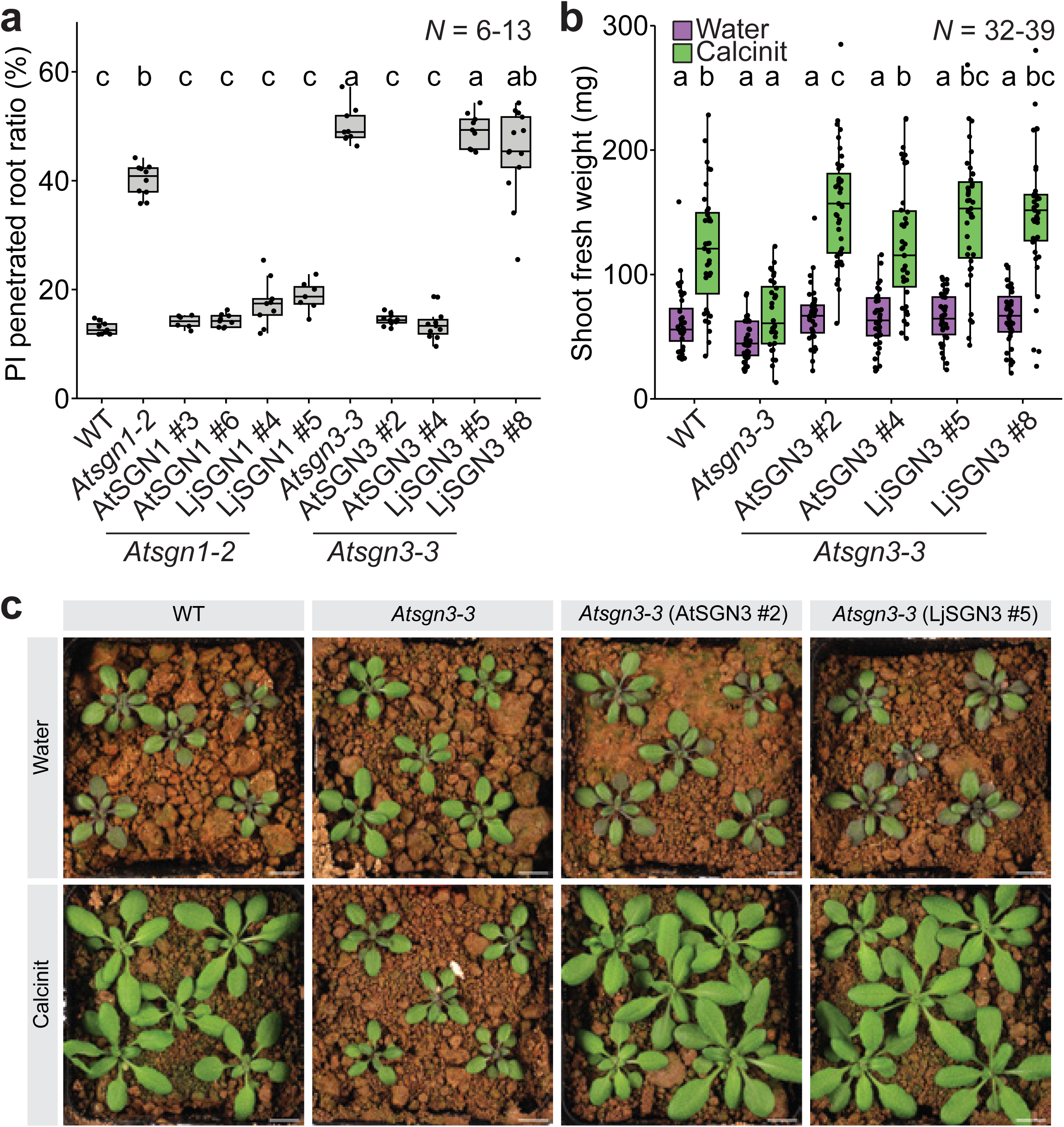
SGN3 mediates shoot nitrogen response independently of root barrier integrity under agricultural soil conditions. a, Proportion of roots exhibiting propidium iodide (PI) penetration in 6-day-old seedlings. **b**, Shoot fresh weight of plants grown in Cologne agricultural soil (CAS) under water-treated or nitrogen-supplemented (Calcinit) conditions **c**, Representative images of 28-day-old plants grown in CAS under water-treated and Calcinit conditions. Different letters in (**a**) and (**b**) indicate statistically significant differences determined by one-way ANOVA followed by Tukey’s multiple-comparison test and Sidak’s multiple-comparison test, respectively (*P* < 0.05). In box plots, the center line indicates the median, box limits represent the 25th and 75th percentiles (IQR), and whiskers extend to the most extreme data points within 1.5 × IQR. Individual values are shown as scattered jittered dots. The number of biological replicates is indicated in the graphs. Scale bar, 1 cm.

To determine how this uncoupling is reflected at the transcriptional level, we performed comparative transcriptome profiling of shoots and roots across nitrogen availability regimes. These analyses were performed under standard agar and CAS conditions, which we used as complementary growth contexts rather than as a single-factor environmental test. Nitrogen supplementation induced extensive transcriptional reprogramming in both conditions (Fig. 3a,b). However, the magnitude and composition of these responses differed markedly between agar and CAS conditions, with functional enrichment analyses revealing distinct biological processes (Extended Data Fig. 4a,b). Whereas SGN3-dependent responses were relatively limited under agar conditions, they expanded markedly under CAS conditions (Fig. 3b) and affected a substantially larger set of canonical nitrogen-responsive genes (Fig. 3c and Extended Data Fig. 4c). Consistent with this condition-associated expansion, SGN3-dependent gene sets showed only modest overlap between agar and CAS conditions, as quantified by low Jaccard similarity (shoot: 0.025; root: 0.058). This condition-associated divergence was also evident at the genome-wide level, where SGN3-dependent nitrogen effects differed markedly between agar and CAS conditions (Extended Data Fig. 4d). In roots, SGN3-dependent nitrogen effects were largely uncorrelated between conditions, whereas in shoots they were strongly negatively correlated, indicating a systematic reversal of gene responses supported by both Spearman and Pearson correlations (Extended Data Fig. 4d). Consistent with this pattern, many genes changed the direction of their response between conditions, particularly in shoots (switch fraction = 0.668 in shoots vs 0.461 in roots). Despite this condition-associated divergence, both AtSGN3 and LjSGN3 restored SGN3-dependent nitrogen responses across conditions and tissues in the *Atsgn3-3* mutant, supporting conservation of the nitrogen-signaling output despite their divergent ability to support CS formation (Extended Data Fig. 4e). Together, these findings reveal a conserved SGN3-dependent nitrogen-signaling branch that is separable from CS formation, becomes most apparent under CAS growth conditions, and undergoes extensive condition- and tissue-dependent rewiring at the level of downstream transcriptional outputs.

**Fig. 3.**
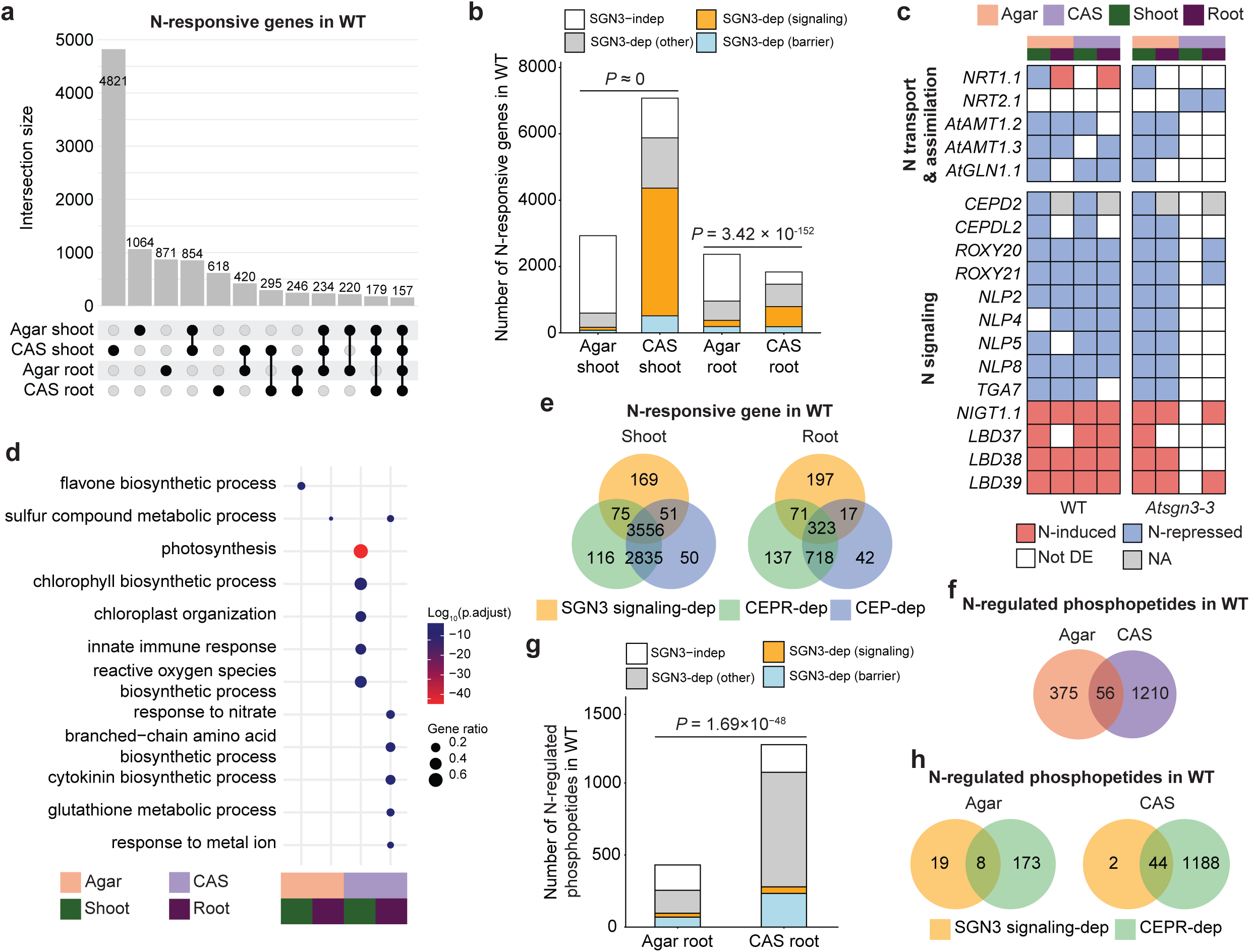
SGN3-dependent nitrogen branch is coupled to CEP–CEPR signaling. **a**, Upset plot showing the overlap among nitrogen-responsive genes (high nitrogen versus low nitrogen; FDR-adjusted *P* < 0.05, absolute fold change > 2) identified in different tissues of wild type (WT) plants grown on agar or in Cologne agricultural soil (CAS). First 12 intersections are shown here (Full intersections data in Source data). **b**, Distribution of gene categories among nitrogen-responsive genes across tissues and growth conditions in WT. Fisher’s exact test was used to compare the proportion of SGN3-dependent genes between agar and CAS conditions in shoots and roots separately. **c**, Heatmap showing the expression patterns of canonical nitrogen-responsive genes in different tissues of WT and *Atsgn3-3* mutant plants grown on agar or in CAS. Red, induced; blue, repressed; white, not differentially expressed (DE); grey, not available due to low expression level (NA). **d**, Dotplot of Gene Ontology terms enriched among SGN3 signaling dependent nitrogen-responsive genes in different tissues and growth conditions of WT plants. **e**, Venn diagram showing the overlap among SGN3-dependent, CEPR-dependent and CEP-dependent nitrogen-responsive genes in WT shoots (left) and roots (right). **f**, Venn diagram showing the overlap among phosphopeptide changes detected in WT roots grown on agar or in CAS. **g**, Distribution of nitrogen-regulated phosphopeptides across different categories in WT roots grown under agar or CAS conditions. Fisher’s exact test was used to compare the proportion of SGN3-dependent phosphopeptide changes between agar and CAS conditions. **h**, Venn diagram showing the overlap between SGN3-signaling-dependent and CEPR-dependent phosphopeptide changes detected in WT roots grown on agar (left) or in CAS (right).

### SGN3-dependent nitrogen branch is coupled to CEP–CEPR signaling

The ability of LjSGN3 to restore nitrogen responses without restoring CS formation allowed us to distinguish SGN3-dependent signaling-associated outputs from barrier-associated responses. We therefore classified nitrogen-responsive genes according to their complementation behavior in *Atsgn3-3* in each tissue and condition. Genes whose nitrogen responsiveness was restored by both AtSGN3 and LjSGN3 were defined as SGN3 signaling-associated outputs, whereas genes restored only by AtSGN3 were classified as barrier-associated outputs (Extended Data Fig. 5). This operational classification does not imply that signaling- and barrier-associated genes are direct SGN3 targets, but uses the differential complementation behavior of AtSGN3 and LjSGN3 to separate outputs associated with nitrogen signaling from those associated with restoration of CS function. This separation revealed that signaling- and barrier-associated nitrogen responses were largely tissue- and condition-specific, with limited overlap at both gene and functional levels (Extended Data Fig. 6). Under CAS conditions, for example, signaling-associated nitrogen-responsive genes in shoots were enriched for photosynthesis and immune-related processes, whereas those in roots were enriched for nitrate responses and amino acid biosynthesis, consistent with engagement of core nitrogen assimilation pathways (Fig. 3d). Systemic nitrogen-demand signaling involves the CEP–CEPR module, in which nitrogen-starvation-induced root-derived CEP peptides are perceived by CEPR1/CEPR2 in the shoot, leading to induction of shoot-derived CEP DOWNSTREAM (CEPD) signals that promote nitrate uptake in roots^16–18^. Reduced nitrogen-responsive expression of CEPR downstream genes, including *CEPD2* and *CEPDL2*, in *Atsgn3-3* shoots under CAS conditions prompted us to test whether the SGN3-dependent nitrogen branch is connected to CEP–CEPR signaling (Fig. 3c). Given the large number of *CEP* family members in *Arabidopsis*, we generated a knockout line targeting all 12 group I *CEP* genes, a subgroup previously associated with nitrogen-starvation responses (CEP1-CEP12, Extended Data Fig. 7)^19^. Under low-nitrogen CAS conditions, both the *CEP* multi-gene knockout (*Atcep* 12× KO) and the CEP receptor double mutant *Atcepr1-3 cepr2-4* phenocopied the nitrogen-response defects of *Atsgn3-3*, while maintaining a functional CS (Extended Data Fig. 8a-d). This supports a role for the CEP–CEPR pathway in regulating nitrogen-dependent growth under soil conditions. Consistently, most nitrogen-responsive genes in WT depended on CEP and/or CEPR function (shoots: 94.4%, roots: 71.2%, Extended Data Fig. 8e), and the majority of the SGN3 signaling-associated outputs overlapped with this subset (shoots: 95.6%, roots: 67.5%; Fig. 3e). To determine whether the separation between SGN3 signaling- and barrier-associated outputs is also reflected at the post-translational level, we performed a phosphoproteomic profiling of roots across the same conditions. Nitrogen availability triggered extensive phosphorylation changes, with substantially more nitrogen-responsive phosphopeptides detected under CAS than agar conditions (Fisher’s exact test, *P* = 1.76×10^−136^), and limited overlap between conditions (Fig. 3f). SGN3 influenced both the extent and structure of these responses, with a more prominent contribution under CAS conditions (Fig. 3g, Extended Data Fig. 9a). This is consistent with the condition-associated difference observed at the transcriptional level (Fig. 3b). Importantly, signaling- and barrier-associated phosphorylation changes remained largely distinct (Extended Data Fig. 9b), mirroring the separation observed at the transcriptional level (Extended Data Fig. 6). CEPR also contributed more strongly to nitrogen-responsive phosphoregulation under CAS conditions, accounting for 42.0% of changes on agar, but 97.3% in CAS (Extended Data Fig. 9c). Moreover, the majority of the SGN3 signaling-associated phosphorylation changes in CAS were CEPR dependent (agar: 29.6%, CAS: 95.6%; Fig. 3h). Together, these findings functionally couple the SGN3-dependent nitrogen branch to CEP–CEPR signaling at both transcriptional and post-translational levels, particularly under CAS conditions.

### Differential ligand perception separates SGN3 signaling outputs

We next investigated whether differences in receptor-ligand perception could explain the separation between SGN3-dependent nitrogen signaling and barrier formation. Because CS-defective *Atcif1 cif2* mutants fail to mount normal nitrogen responses under CAS conditions^10^, the CIF–SGN3 module likely contributes to both barrier-associated and nitrogen-signaling outputs. Although nitrogen-response complementation assays with exogenous peptides are not feasible under CAS conditions, peptide-dose agar assays in the *Atcif1 cif2* double mutant showed that low concentrations of AtCIF2_sTyr2_ were sufficient to restore CS formation, whereas higher concentrations of LjCIF2_sTyr2_ were required to achieve comparable effects (Extended Data Fig. 10a). A similar pattern was observed in ectopic suberization assays, where native receptor-ligand combinations induced stronger oversuberization than non-native combinations (Extended Data Fig. 10b).

To test whether the CS branch of SGN3 outputs depends on effective ligand perception, we examined receptor variants with altered ligand-binding properties. Replacing the LjSGN3 ectodomain with that of AtSGN3 restored CS formation and ligand responsiveness, despite retaining the *Lotus* cytosolic kinase domain (Fig. 4a-c), indicating that ligand recognition is a key determinant of output selection. Progressive introduction of AtSGN3 CIF-binding residues into LjSGN3 enhanced its ability to support CS formation and restore CS-domain integrity in *Atsgn3-3* (Fig. 4d,e and Extended Data Fig. 11). Conversely, AtSGN3 variants with reduced ligand-binding capacity partially restored CS integrity^20^ but substantially rescued nitrogen responses in *Atsgn3-3* under CAS conditions (Fig.4f and Extended Data Fig. 12). Thus, manipulations that reduce effective ligand perception preferentially compromise CS formation while preserving nitrogen signaling, indicating that nitrogen signaling can be retained under receptor-perception states that are insufficient for full CS formation.

**Fig. 4.**
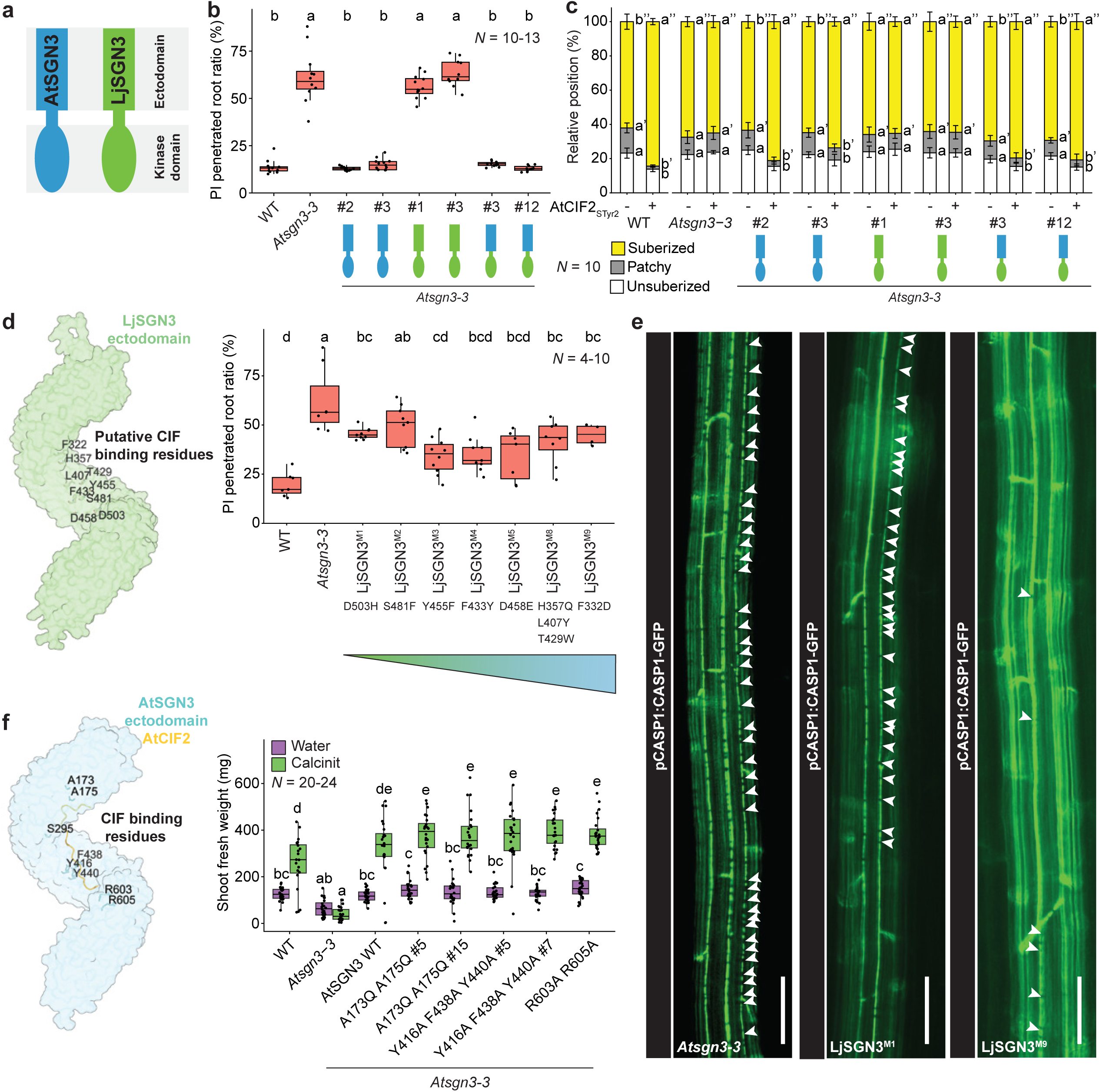
Different SGN3 ligand perceptions underpin Casparian strip formation and shoot nitrogen response. a, Schematic illustration of the ectodomain and kinase domain of SGN3 from *Arabidopsis* (blue) and *Lotus* (green). **b,** Proportion of roots exhibiting propidium iodide (PI) penetration in 7-day-old seedlings. **c**, Suberization patterns of 7-day-old *Atsgn3-3* complementation lines following 2 days of treatment with water or 10 nM AtCIF2. Different letters indicate statistically significant differences determined by one-way ANOVA followed by Tukey’s multiple-comparison test (P < 0.05) within the same genotype. **d**, Left: AlphaFold predicted structure of the LjSGN3 ectodomain. Amino acids associated with CIF2 binding that differ from their counterparts in AtSGN3 are highlighted and subjected to mutagenesis. Right, proportion of roots exhibiting PI penetration in 9-day-old seedlings. Mutations were introduced progressively, with residues accumulated from LjSGN3^M1^ to LjSGN3^M9^. **e**, Representative images of CASP1-GFP signals in 9-day-old *Atsgn3-3* seedling roots and *Atsgn3-3* complemented with mutant LjSGN3 variants. Arrowheads indicate discontinuities in the Casparian strip domains visualized by CASP1-GFP signals. **f**, Left: published structure of the AtSGN3 ectodomain (Protein Data Bank under ID code 6S6Q). Amino acids previously shown to be involved in CIF2 binding are highlighted. Right: Shoot fresh weight of plants grown in Cologne agricultural soil (CAS) under water-treated or nitrogen-supplemented (Calcinit) conditions. Different letters in (**b**-**d**) indicate statistically significant differences determined by one-way ANOVA followed by Tukey’s multiple-comparison test (*P* < 0.05). Different letters in (**f**) indicate statistically significant differences determined by one-way ANOVA followed by Sidak’s multiple-comparison test (*P* < 0.05). In bar charts, error bars denote standard deviation. In box plots, the center line indicates the median, box limits represent the 25th and 75th percentiles (IQR), and whiskers extend to the most extreme data points within 1.5 × IQR. Individual values are shown as scattered jittered dots. The number of biological replicates is indicated in the graphs. Scale bar, 50 µm.

We next asked whether co-receptor function also contributes to SGN3 output selection. Members of the SOMATIC EMBRYOGENESIS RECEPTOR KINASE (SERK) family act as ligand-induced co-receptors for diverse receptor kinases, including SGN3, and can modulate signaling strength and specificity^20,21^. Mutation of *SERK1* alone as well as higher-order *serk1 serk2 serk4* and *serk3 serk4 serk5* combinations impaired nitrogen responses while maintaining normal CS formation, whereas *bak1-5*, a semi-dominant *SERK3* mutation that can interfere with SERK-dependent receptor signaling^22^, affected both nitrogen responses and CS formation (Extended Data Fig. 13). Thus, co-receptor composition contributes to SGN3 output selection, potentially by altering effective receptor activation state or downstream effector engagement.

## Discussion

Our results establish SGN3 as a signaling integrator that links ligand perception state to distinct biological outputs. Although SGN3 has been primarily associated with CS formation, we show that it also controls systemic nitrogen responses and that these functions are genetically separable. Cross-species complementation uncouples nitrogen signaling from barrier formation, while transcriptomic and phosphoproteomic analyses reveal largely distinct signaling- and barrier-associated programs. Notably, SGN3-dependent nitrogen responses are most prominent under CAS growth conditions, indicating that this branch represents a physiologically relevant output that is comparatively weak in standard agar-based assays.

Comparison with *Lotus* reveals that the connection between endodermal barrier pathways and systemic nitrogen signaling is conserved, but organized differently. In *Lotus*, SGN activation relies on a MYB36-dependent competence state, consistent with observations in rice, where expression of both *SGN1* and *SGN3*, as well as SGN3 signaling, is largely MYB36-dependent^23^. SGN3 signaling cannot be triggered by CIF peptides alone in *Lotus*, despite the ability of *Lotus* CIF peptides to activate SGN3 outputs in *Arabidopsis*. Thus, ligand perception is not universally sufficient for SGN pathway activation. Instead, evolutionary changes in pathway deployment can place SGN signaling under different developmental or transcriptional control while preserving its broader role in linking endodermal status to systemic physiology. In support of this, we recently showed that CS is associated with systemic nitrogen signaling to initiate root nodule formation in *Lotus*^11^. Therefore, one possible explanation is that legumes have rewired barrier-associated signaling in the context of nodulation. Nodule formation requires extensive reprogramming of cortical and endodermal tissues and is tightly integrated with nitrogen signaling^24,25^. It is therefore tempting to speculate that placing SGN activation under a MYB36-dependent competence state could prevent inappropriate barrier-surveillance activation during symbiotic re-differentiation. This idea remains speculative, but highlights how conserved receptor modules may be redeployed according to lineage-specific developmental demands.

Mechanistically, our data support a model in which effective SGN3 ligand perception biases output selection. Receptor-ligand comparisons, engineered SGN3 variants and SERK analysis indicate that systemic nitrogen signaling can be retained under receptor-perception states that are insufficient for full CS formation (Extended Data Fig. 14). This provides a simple explanation for how LjSGN3 can restore nitrogen responses without restoring CS function, and how reduced-perception AtSGN3 variants can separate these outputs. Our data are consistent with an amplitude-based threshold model, in which nitrogen signaling requires a lower level of SGN3 activation than barrier formation. However, they do not distinguish this from a branched-output model, in which different receptor-perception states engage qualitatively distinct downstream effectors. The largely non-overlapping signaling- and barrier-associated transcriptomic and phosphoproteomic programs are compatible with such a branched architecture. Future quantitative receptor-activation reporters and direct binding measurements across receptor variants will be required to resolve the molecular implementation of SGN3 output selection. This SGN3-dependent nitrogen branch is closely coupled to the CEP–CEPR module at both transcriptional and post-translational levels, placing SGN3 in close functional connection with canonical systemic nitrogen-demand signaling.

Together, these findings identify ligand perception state as a mechanism for receptor output specificity. Rather than acting as a binary barrier-integrity switch, SGN3 functions as a tunable receptor module in which ligand structure, receptor ectodomain properties and co-receptor engagement bias the biological output. This framework explains how a shared receptor module can selectively engage local barrier formation or systemic nutrient signaling, allowing plants to coordinate root barrier state with whole-plant physiology across growth contexts.

## Methods

### Plant materials and growth conditions

*Lotus japonicus* (*Lotus*) accession Gifu B-129 (Gifu) was used as the wild type (WT), unless otherwise specified. *Lotus* seeds were scarified with sandpaper and surface-sterilized with 1% sodium hypochlorite for 10 min, rinsed six times with sterile Milli-Q water, and incubated in sterile Milli-Q water at room temperature for 1 h with shaking. Seeds were stratified on wet filter paper in Petri dishes (90 mm diameter) at 4 °C in darkness for 2–3 d, then germinated at 21 °C in darkness for 2 d. Seedlings were transferred to square plates (12 × 12 cm) containing 40 ml of ¼ B&D medium supplemented with 0.5 mM KNO₃ and grown at 21 °C under a 16 h light/8 h dark photoperiod. For *Lotus* growth, ¼ B&D medium was always supplemented with 0.5 mM KNO_3_.

*Arabidopsis thaliana* (*Arabidopsis*) Col-0 plants expressing the *pAtCASP1:AtCASP1-GFP* construct were used as the WT control for *Atsgn3-3* complementation lines (with LjSGN3, chimeric LjSGN3, and LjSGN3 mutant variants) and for the transcriptome and phosphoproteomic analysis of *Atcep* 12 × KO and *Atcepr1-3cepr2-4* KO^26^. Unmodified Col-0 plants were used as the WT control for *Atsgn3-3* complementation lines with AtSGN3 variant, phenotyping of *Atcep* 12 × KO and *Atcepr1-3 cepr2-4*. *Arabidopsis* seeds were surface-sterilized in 70% ethanol containing 0.05% Triton X-100 for 5 min, rinsed briefly with 100% ethanol, and air-dried. Seeds were sown on ½ Murashige and Skoog (MS) medium, stratified at 4 °C in darkness for 2–3 d, and grown at 21 °C under a 16 h light/8 h dark photoperiod. For transcriptomic and phosphoproteomic analyses under low-and high-nitrogen growth conditions, seeds were sown on nylon meshes (SEFAR NITEX 03-100/44, cut into 11 cm × 11 cm squares) placed on top of nitrogen-free MS medium (MSP21, Caisson Labs). The medium was supplemented with 10 mM MES and 1% Duchefa plant agar (pH 5.8), and further amended with either 0 mM KNO₃ (low nitrogen) or 30 mM KNO₃ (high nitrogen).

### CIF2 peptide treatments

For agar-based treatments, 6-day-old *Lotus* seedlings were transferred to ¼ B&D agar medium supplemented with either 500 nM CIF2 peptide or H₂O (mock) and grown for 2 d. For hydroponic treatments, 2-day-old germinated seedlings were first transferred to sterile WECK jars containing glass beads (5 mm diameter) and ¼ B&D liquid medium. At 6 days of age, seedlings were treated with either 500 nM peptide or H_2_O (mock) by refreshing the liquid medium and then grown for an additional 2 days. For TAMRA-labelled AtCIF2 treatment, 8-day-old *Lotus* seedlings was treated with either 500 nM TAMRA-labelled AtCIF2 peptide or H_2_O (mock) in a coverslip-mounted glass chamber under darkness for 2 h.

For *Arabidopsis*, 5-day-old seedlings grown on ½ MS medium were transferred to ½ MS agar plates supplemented with 10–100 mM CIF2 peptide or H₂O (mock) and grown for 2 d.

### Generation and genotyping of mutants

*Ljmyb36-2 × sgn3-2* double mutants were generated by crossing emasculated *Ljmyb36-2* plants with *Ljsgn3-2* (ref ^11^). Homozygous double mutants were identified in the F₂ population. LORE1 insertion mutants were genotyped using CTAB-based DNA extraction. Wild type sibling lines corresponding to each LORE1 allele were identified and used as controls where indicated. The CRISPR-Cas9-generated *Atcep* 12*×* knockout mutant was generated in the previously described *Atcep 6×* knockout mutant background, which carries loss-of-function mutations in *CEP1*-*CEP4*, *CEP6* and *CEP9* (Ref. ^26^). Putative edited plants were screened by PCR using gene-specific primers, and all mutations were confirmed by Sanger sequencing of the corresponding amplicons. Primer sequences together with mutant lines used in this study are listed in Supplementary Table 1.

### Plasmid construction and plant transformation

Constructs for *Lotus* transformation were generated using the Golden Gate cloning system^27^. The promoter region of *LjSCR* (3137 bp) was cloned into pICH41295. Coding sequences (CDSs) of *LjSGN1* (1281 bp) and *LjSGN3* (3819 bp) were cloned into pICH41308, whereas the 3′ untranslated regions (UTRs) of *LjSGN1* (595 bp) and *LjSGN3* (530 bp) were cloned into pICH41276. Level 1 constructs were assembled from level 0 modules using pICH47751 or pICH47761 as acceptor vectors. The constructs *pLjSCR:LjSGN1:act2Ter* (pICH44300) and *pLjSCR:GUS:act2Ter* (pICH75111) were assembled in pICH47751. The construct *pLjSCR:LjSGN3:act2Ter* was assembled in pICH47761 and used for coexpression with *LjSGN1*. For selection and visualization of transgenic roots, *pAtUBQ10:NLS-3×mScarlet:nosTer* (pICH41421) was assembled in pICH47732, and *pNos:NPTII:OcsTer* (pICSL70004) was cloned into pICH47742. Level 2 constructs were assembled in pAGM4673 and introduced into *Agrobacterium rhizogenes* strain MSU440 by electroporation.

Constructs for *Arabidopsis* transformation were generated using the Gateway cloning system (Invitrogen). CDSs of *LjSGN1* and *LjSGN3* were first cloned into pDONR221. For generation of a chimeric *SGN3* construct, the N-terminal region of *AtSGN3* (1–1260 bp) was fused to the C-terminal region of *LjSGN3* (2659–3816 bp) by PCR. For chimeric SGN3 variants, the coding sequence of mTurquoise2 was fused in-frame to the C-terminus of *AtSGN3*, *LjSGN3*, or the chimeric *AtSGN3/LjSGN3* construct (without an internal stop codon) and cloned into pDONR221. For the point-mutant variants of LjSGN3, the CDSs were synthesized and cloned into pDONR221. Mutated residues were chosen from the resolved structure of the AtSGN3 ectodomain^20^, focusing on ligand-contacting positions that are not conserved in LjSGN3. A series of variants, designated LjSGN3^M1^, LjSGN3^M2^, LjSGN3^M3^, LjSGN3^M4^, LjSGN3^M5^, LjSGN3^M8^, LjSGN3^M9^, was generated by successively introducing point mutations that convert the corresponding LjSGN3 residues to their *Arabidopsis* counterparts. Accordingly, LjSGN3^M1^ carries a single mutation, whereas LjSGN3^M9^ contains nine mutations. Promoter and terminator sequences (*AtSGN1*, *AtSGN3*, *AtSGN1* 3′ UTR, and *AtSGN3* 3′ UTR)^28,29^, together with entry clones, were recombined into the binary vector pED97^30^. The resulting constructs were introduced into *Atsgn1-2* or *Atsgn3-3* mutant backgrounds expressing *pAtCASP1:AtCASP1-GFP*^28,29^ via *Agrobacterium tumefaciens* GV3101 using the floral dip method^31^. Transgenic seeds expressing the FastRed marker were selected using a Zeiss Axio Zoom.V16 stereomicroscope. Independent homozygous T3 lines were used for all analyses, except for Fig. 4d and Extended Data Fig. 11b, where T1 lines were used. Primers used for cloning are listed in Supplementary Table 1.

For generation of the CRISPR–Cas9-mediated *Atcep* 12× knockout mutant, the remaining group I *CEP* genes, *CEP5*, *CEP7*, *CEP8*, *CEP10*, *CEP11* and *CEP12*, were targeted using two guide RNAs per gene. Nineteen-nucleotide target sites were designed using CHOPCHOP (https://chopchop.cbu.uib.no/). Each sgRNA expression cassette comprised the promoter and terminator sequences of AT3G13855, a gene-specific target sequence and a guide RNA scaffold, flanked by BpiI recognition sites. Two sgRNA cassettes per target gene were synthesized commercially (Twist Bioscience, USA). The 12 sgRNA cassettes were assembled into three intermediate guide RNA arrays in a Golden Gate-adapted pUC18-based vector, with each array comprising guides targeting two *CEP* genes (two target sites per gene). The resulting three guide RNA arrays were subsequently assembled by BsaI-mediated Golden Gate cloning together with the FastRed-pRPS5::Cas9 cassette into pICSL4723OD for plant transformation via *Agrobacterium tumefaciens* GV3101 using floral dip method^31,32^. T1 transformants carrying the CRISPR–Cas9 construct were identified by FastRed seed fluorescence and grown on soil. T2 progeny lacking FastRed fluorescence were selected to identify individuals that had segregated away the CRISPR-Cas9 transgene. These lines were subsequently propagated and genotyped until homozygous and transgene-free. Target sequences are listed in Supplementary Table 1.

### Staining procedures and microscopy

Propidium iodide (PI, Sigma-Aldrich, P4170) and Fluorol Yellow 088 (FY; Interchim, FP-1J8050) staining of *Arabidopsis* and *Lotus* roots were performed as previously described^33,34^. Suberization patterns in *Arabidopsis* roots were determined as described previously^35^. Suberization patterns in *Lotus* roots were defined as follows: the “patchy zone” was identified at the position of the first suberized endodermal cell, typically associated with phloem poles. The “continuous zone” was defined as the region in which endodermal cells associated with phloem poles were continuously suberized (see Extended Data Figure 1a).

Confocal imaging of PI penetration (*Arabidopsis* and *Lotus*) and FY staining (*Lotus*) was performed using a Zeiss LSM 980 confocal microscope. PI was excited at 514 nm, and emission was detected at 600–650 nm; FY was excited at 488 nm, and emission was detected at 500–550 nm. FY staining in *Arabidopsis* roots was imaged using a Zeiss Axio Zoom.V16 stereomicroscope equipped with a Zeiss Axiocam 705 mono or color camera.

### Hairy root transformation

Transient root transformation in *Lotus* was performed using the hairy root method^36^. 7-day-old seedlings grown on Gamborg’s B5 agar plates were excised, and hypocotyls were immersed in a suspension of *Agrobacterium rhizogenes* strain MSU440 carrying the indicated constructs. Infected hypocotyls were placed on Gamborg’s B5 agar plates and incubated at 21 °C in darkness for 2 d, followed by incubation at 21 °C under a 16-h light/8-h dark photoperiod for an additional 3 d. Hypocotyls were then transferred to Gamborg’s B5 agar plates supplemented with cefotaxime (300 µg mL⁻¹) to eliminate *A. rhizogenes* and neomycin (25 µg mL⁻¹) to suppress the growth of non-transformed roots. Composite plants used for axenic root development assays were transferred to ¼ B&D agar plates and grown for 7 d. For β-glucuronidase (GUS) staining, roots expressing promoter:GUS constructs were incubated in staining buffer containing 0.05% (w/v) X-Gluc (5-bromo-4-chloro-3-indolyl β-D-glucuronide), 100 mM phosphate buffer (pH 7.0), 0.5 mM EDTA (pH 7.0), 0.5 mM potassium ferricyanide [K₃Fe(CN)₆], and 0.5 mM potassium ferrocyanide [K₄Fe(CN)₆]. Samples were vacuum-infiltrated for 15 min at room temperature and incubated at 37 °C for approximately 2 h. Samples were subsequently washed three times with 100 mM phosphate buffer (pH 7.0) prior to imaging. Whole-root images were acquired using a Zeiss Axio Zoom.V16 stereomicroscope equipped with a Zeiss Axiocam 705 color camera.

### Growth on Cologne agricultural soil

For Cologne agricultural soil (CAS)-based growth assays, 7-day-old *Arabidopsis* seedlings were transferred to square pots (9 × 9 cm) containing CAS. Pots were placed on a capillary mat and irrigated from below with either tap water or 7% Calcinit™ solution (v/v; 0.651 g l⁻¹; containing 14.4% nitrate, 1% ammonium, and 26% calcium oxide; Yara, Germany). Plants were grown at 21 °C under a 16 h light/8 h dark photoperiod. Plants were imaged and harvested after 2-3 weeks on CAS.

### Peptides

AtCIF2_sTyr2_ peptides [DY(sulphated)GHSSPKPKLVRPPFKLIPN] were synthesized by Pepmic Co., Ltd (Suzhou, China). LjCIF2_sTyr2_ peptides [DY(sulfated)GRYDPTPKLSKPPFKLIPN] and LjCIF2_sTyr2Tyr5_ peptides [DY(sulfated)GRY(sulfated)DPTPKLSKPPFKLIPN] were synthesized by GL Biochem Ltd (Shanghai, China).

### Anthocyanin content measurement

Measurement of anthocyanin content in *Arabidopsis* rosettes grown under CAS conditions was performed as previously described^37^. Absorbances at 530 and 637 nm were measured using a Tecan Infinite 200 PRO plate reader.

### Transcriptomics

For transcriptomic analysis of *Lotus* roots, whole roots from 8-day-old seedlings grown on ¼ BD agar plates were harvested, with approximately 10 roots pooled per biological replicate. Samples were immediately frozen in liquid nitrogen. For transcriptomic analysis of *Arabidopsis* plants grown on agar medium, 7-day-old seedlings were germinated and grown on the indicated medium. Shoots and roots were excised and harvested separately. For transcriptomic analysis of *Arabidopsis* plants grown on CAS, 21-day-old plants were prepared as described above. Whole plants were gently removed from the soil under running tap water, and shoots and roots were briefly dried on paper towels and harvested separately.

RNA was extracted using a TRIzol (Invitrogen)-adapted ReliaPrep RNA extraction kit (Promega), as previously described^30^. RNA quality was determined using a Bioanalyzer 2100 system (Agilent Technologies). Library preparation and paired-end 150 bp sequencing were conducted by Novogene (Cambridge, UK). Approximately 40 million raw reads were generated per sample. RNA-seq raw reads of Col-0, *Atmyb36-2*, *Atsgn3-3 and Atmyb36-2xsgn3-3* roots were acquired from Reyt et al.^13^. All raw reads were preprocessed using fastp (v0.22.0)^38^; filtered high-quality reads were mapped to *Arabidopsis* TAIR10 reference genome with Araport 11 annotation (Phytozome genome ID: 447) or *Lotus* Gifu v1.2 genome assembly^39^ with Gifu v1.3 annotation (https://lotus.au.dk) using HISAT2 (v2.2.1)^40^, and counted using featureCounts from the Subread package (v2.0.1).^41^ All statistical analyses were performed using R (https://www.R-project.org/).

For comparative analyses of *Lotus* and *Arabidopsis* Casparian strip mutants, raw read counts were further analyzed as previously described^10,11^. Differentially expressed genes (DEGs) were defined as those with a false discovery rate (FDR) < 0.05 and an absolute fold change > 2, and were used for downstream analyses. Gene Ontology (GO) enrichment analysis was performed using the topGO package (https://github.com/federicomarini/topGO) in R, as clusterProfiler does not support *Lotus* gene annotations. Gene-to-GO annotations were derived from eggNOG^42^ (*Lotus*) and TAIR (*Arabidopsis*). Enrichment was assessed using Fisher’s exact test with the *elim* algorithms, GO terms with *elim P* ≤ 0.05 were considered significantly enriched.

For *Atsgn3-3* complementation lines, raw count data were filtered to remove lowly expressed genes using the filterByExpr function from the edgeR package, followed by normalization with the TMM (trimmed mean of M-values) method; differential expression analysis was performed using the limma-voom framework with a linear model fitted with genotype, treatment, and their interaction (∼ Genotype × Treatment), and empirical Bayes moderation was applied to improve variance estimation. Genotype-specific nitrogen responses were assessed using linear contrasts, allowing the estimation of HighN versus LowN effects within each genotype, including WT, *Atsgn3-3*, *AtSGN3*, *LjSGN3*, *Atcepr1cepr2*, and *Atcep* 12× KO. Nitrogen-responsive genes in WT were defined as those with an FDR < 0.05 and an absolute log₂ fold change > 1 (|log₂FC| > 1); these WT-responsive genes were further categorized into functional groups based on their response patterns across genotypes, including SGN3-dependent/independent (genes that lost or maintained their N response in *sgn3*, respectively), CEPR-dependent/independent (lost or maintained in *Atcepr1cepr2*), CEP-dependent/independent (lost or maintained in *Atcep* 12× KO), signaling-dependent (lost in *Atsgn3-3* but restored in both *AtSGN3* and *LjSGN3*), and CS-dependent (lost in *Atsgn3-3* but restored only in *AtSGN3*, not *LjSGN3*).

Overlap between gene sets identified under different growth conditions was quantified using Jaccard similarity and assessed for enrichment using Fisher’s exact tests based on the expressed-gene universe. To control for differences in gene-set size, overlap robustness was additionally evaluated by power-matched downsampling. Gene lists for each category were extracted for downstream analyses. GO enrichment analysis was performed using the clusterProfiler^43^ package with the *Arabidopsis* annotation database (org.At.tair.db). Enrichment was assessed in the biological process category using a background set of all expressed genes. Results were adjusted for multiple testing using the Benjamini–Hochberg method, and enriched terms with adjusted *P*-value lower than 0.05 were visualized using dot plots. Principal component analysis (PCA) of whole transcripts after filtering was performed using prcomp function in R and visualized using ggplot2 package^44^. Full gene lists and GO term enrichment results of all transcriptome analyses are listed in Supplementary Table 2.

### Proteomics

#### Preparation of phospho-enriched samples from soil- and agar-grown root tissue

For proteomics analysis of *Arabidopsis* plants grown on agar, materials were prepared as described above. A volume of roughly 75-100 µL of Arabidopsis seeds was sterilized and placed in three rows onto the nylon mesh. All seedlings of one plate were pooled to generate one replicate. For proteomics analysis of *Arabidopsis* plants grown on CAS, materials were prepared as described above. The frozen tissue was ground with two 2.5 mm steel beads in a Retsch mill, then 250-600 µL (depending on amount of material and absorption) extraction buffer (8M urea, 20 µL/mL Phosphatase Inhibitor Cocktail 2 (Sigma, P5726-5ML), 20 µL/mL Phosphatase Inhibitor Cocktail 3 (Sigma, P0044-5ML),5 mM DTT) was added and samples were incubated for 30 min with shaking, after which cell debris was removed by centrifugation. Samples were alkylated with CAA (550 mM stock, 14 mM final), the reaction was quenched with DTT (5 mM final). Next, samples were diluted to 4M urea with 100 mM Tris-HCl pH 8.5, 1 mM CaCl_2_ and digested with 5 µg LysC (stock: 1 µg/µL Lys-C (Merck) in H_2_O) for 3 h at RT. Subsequently, samples were diluted with 100 mM Tris-HCl pH 8.5, 1 mM CaCl_2_ to 1M urea, then 5 µg trypsin (stock: 1 µg/µL in 1 mM HCl,) was added and samples were mixed and incubated overnight at 37 °C. After incubation, samples were acidified with TFA to 0.5% final concentration and samples were desalted using C18 SepPaks (1cc cartridge, 100 mg (WAT023590)). In brief, SepPaks were conditioned using methanol (1 mL), buffer B (80% acetonitrile, 0.1% TFA) (1 mL) and buffer A (0.1% TFA) (2 mL). Samples were loaded by gravity flow, washed with buffer A (1 x 1 mL, 1x 2 mL) and eluted with buffer B (2 x 400 µL). 40 µL of eluates were used for peptide measurement and total proteome control.

For phosphopeptide enrichment by metal-oxide chromatography (MOC) (adapted from^45^) the remaining samples were evaporated to a sample volume of 50 µL and diluted with sample buffer (2 mL AcN, 820 µL lactic acid (LA), 2.5 µL TFA / 80% ACN, 0.1% TFA, 300mg/ml LA, final concentrations) (282 µL). Since all samples showed a small pellet after dilution, the samples were centrifuged briefly and only the supernatant was used for phosphopeptide enrichment. MOC tips were prepared by loading a slurry of 3mg/sample TiO_2_ beads (Titansphere TiO_2_ beads 10 µm (GL Science Inc, Japan, Cat. No. 5020-75010)) in 100 µL MeOH onto a C2micro column and centrifugation for 5 min at 1500g. Tips were washed with centrifugation at 1500g for 5 min using 75µL of solution B (80% acetonitrile, 0.1% TFA) and 75 µL of solution C (300 mg/mL LA in solution B). To simplify the processing, samples tips were fitted onto a 96/500 µL deep well plate (Protein LoBind, (Eppendorf Cat. No. 0030504100). After washing MOC tips were transferred to a fresh plate, samples were loaded onto the equilibrated tips and centrifuged for 10 min at 1000g. The flow through was reloaded onto the tips and centrifugation was repeated. Tips were washed with centrifugation at 1500g for 5 min using 75 µL of solution C and 3x 75µL of solution B. For the elution of the enriched phosphopeptides the tips were transferred to a fresh 96/500 µL deep well plate containing 100 µL/well of acidification buffer (20% phosphoric acid). Peptides were eluted first with 50 µL elution buffer 1 (5% NH_4_OH) and centrifugation for 5 min at 800g, then with 50 µL of elution buffer 2 (10% piperidine) and centrifugation for 5 min at 800g. Next, the samples were desalted using StageTips with C18 Empore disk membranes (3 M)^46^, dried in a vacuum evaporator, and dissolved in 10 µL 2% ACN, 0.1% TFA (A* buffer) for MS analysis.

#### LC-MS/MS data acquisition

Samples were analyzed using an Ultimate 3000 RSLC nano (Thermo Fisher) coupled to an Orbitrap Exploris 480 mass spectrometer equipped with a FAIMS Pro interface for Field asymmetric ion mobility separation (Thermo Fisher). Peptides were pre-concentrated on an Acclaim PepMap 100 pre-column (75 µM x 2 cm, C18, 3 µM, 100 Å, Thermo Fisher) using the loading pump and buffer A** (water, 0.1% TFA) with a flow of 7 µl/min for 5 min. Peptides were separated on 16 cm frit-less silica emitters (New Objective, 75 µm inner diameter), packed in-house with reversed-phase ReproSil-Pur C18 AQ 1.9 µm resin (Dr. Maisch). Peptides were loaded on the column and eluted for 130 min using a segmented linear gradient of 5% to 95% solvent B (0 min : 5%B; 0-5 min -> 5%B; 5-65 min -> 20%B; 65-90 min ->35%B; 90-100 min -> 55%; 100-105 min ->95%, 105-115 min ->95%, 115-115.1 min -> 5%, 115.1-130 min ->5%) (solvent A 0% ACN, 0.1% FA; solvent B 80% ACN, 0.1%FA) at a flow rate of 300 nL/min. Mass spectra were acquired in data-independent acquisition (DIA) mode with field asymmetric ion mobility separation (FAIMS) using two compensation voltages (CV) (-45 and -65). For each CV a full scan was performed for mass range of 300–1650 m/z at a resolution of 120,000 FWHM and a normalized AGC target of 300%. Inject time was set to 20 ms. The full scan was followed by 30 variable window DIA scans from 300-1650 m/z, windows overlapped by 0.5 m/z. HCD fragmentation was performed at stepped collision energies of 25, 27.5 and 30%. MS/MS spectra were acquired with normalized ACG target of 1000% at a resolution of 30,000 FWHM, at an automated injection time.

#### Data analysis

Total proteome data was analyzed using Spectronaut software (20.2.250922.92449(soil-grown samples) and 20.5.260227.92449 (agar-grown samples) using DIA+ mode (library-free). Samples were searched against a database from *A. thaliana*(TAIR10_pep_20101214; ftp://ftp.arabidopsis.org/home/tair/Proteins/TAIR10_protein_lists/), trypsin/P specificity was required, otherwise the default settings (BGS phospho PTM Workflow) were used. The output of the Spectronaut search were exported as a customized Pivot Peptide Report containing intensities of each precursor identified. For the downstream analysis the output was filtered for phospho(STY)-modified precursors and the analysis was carried out on phosphopeptide level. Statistical analysis of the intensity values was carried out using Perseus (version 1.6.14.0, http://www.maxquant.org/). First, values were log2 transformed and samples were grouped by condition and divided into subsets. Next, the data of each subset was separated for a mixed imputation processing: hits were filtered for 3 valid values in one of the conditions. Then, the data was separated into two sets: one set containing mostly missing at random (MAR) hits and the other set containing mostly missing not at random (MNAR) hits by filtering the data for 1 valid hit in each group and splitting the resulting matrices^47^. The resulting matrix with at least 1 valid hit in each group is the MAR dataset, the matrix with the hits filtered out is the MNAR dataset. The missing values of the MAR dataset were then imputed using a nearest neighbor approach (KNN, n=4), of the “imputeLCMD” R package (https://CRAN.R-project.org/package=imputeLCMD) integrated into Perseus. The missing values from the MNAR dataset were imputed using the normal distribution imputation featured in Perseus using the default settings (1.8 downshift, separately for each column). After merging of the imputed datasets two-sample Student’s *t*-tests were performed using a permutation-based FDR of 5%. The resulting phosphopeptide-level fold-change tables were imported into R for downstream classification. Phosphopeptides with |log2 fold change| >1 (agar: 30 mM KNO_3_ vs 0 mM KNO_3_; CAS: Calcinit vs water) and FDR < 0.05 in WT were considered nitrogen responsive and were assigned to SGN3-dependent, CEPR-dependent, signaling-dependent, or Casparian strip–dependent categories according to their response patterns in mutant and complementation lines, in a similar manner as the transcriptome analysis. Only phosphopeptides detected across all genotypes were retained for analysis.

### Quantitative RT-PCR

For qRT-PCR on hairy roots, materials were prepared as described above. About five independent axenic root tips (∼2 cm) were sampled as one biological replicate. RNA extraction was performed using a TRIzol (Invitrogen)-adapted ReliaPrep RNA extraction kit (Promega), as described above. cDNA synthesis was performed using iScript™ cDNA Synthesis Kit (BioRad). qRT-PCR was performed in a BioRad CFX Connect Real-Time system in a final volume of 10 μL. Each reaction contained 5 μL of 2 x iQ SYBR Green supermix (Bio-Rad), 2 μL of diluted cDNA (10 times dilution for shoots and roots, 2 times dilution for susceptible zones with water), 1 µL of 2.5 μM forward primer, 1 µL of 2.5 μM reverse primer and 1 µL of water. *LjPP2A* and *LjATPase* were used as the reference housekeeping genes. The thermal cycler conditions were: 95 °C for 2min, 40 cycles of 95°C for 30 s, 60°C for 30 s, and 72°C for 30 s. For the melting curve, conditions were set as: denaturation, 95°C for 10 s; hybridization, 60°C for 5 s; denaturation until 95°C with 0.5°C incrementation. Relative expression values were determined using the 2^-ΔΔCT^ method. qRT-PCR primers used in this study are listed in Supplementary Table 1.

### Phylogenetic Analysis

For *Lotus* and *Medicago truncatula* protein sequences, a local BLASTP was performed against Gifu v1.3 protein and *M. truncatula* A17 r5.0 protein sequences (cut-off Expect (E) threshold < -5). Protein sequences of other species were retrieved from Phytozome v13 by BLASTP using *Arabidopsis* protein sequences (cut-off Expect (E) threshold < - 5). Full-length protein sequences were aligned using MAFFT v7.450^29^ with default parameter settings (auto algorithm; scoring matrix, BLOSUM62; gap opening penalty, 1.53; offset value: 0.123). Curated alignment was used for tree building using FastTree v2.1.11 with the default parameter settings. Branch support analysis was performed using the Shimodaira-Hasegawa test on the three alternate topologies (NNIs) around that split, based on 1000 resamples. Only branches containing genes of interest and their closest homologs were sampled again, aligned and a final phylogeny tree was built as described above. *Lotus* CIF1 and CIF2 protein sequences were not annotated in Gifu v1.3, therefore retrieved from MG20 v3.0 by a local BLASP. A phylogenetic tree on full-length protein sequences was built as above-mentioned, only the regions encoding mature CIF1 and CIF2 peptides were shown in Extended Data Figure 1c.

### Acronyms and accession numbers

All acronyms with accession numbers used in this study are listed in Supplementary Table 5.

### Data availability

RNA-seq raw reads have been deposited at NCBI under PRJNA1370857, PRJNA1370960, PRJNA1085518 and PRJNA1022988. Proteomics data are available via ProteomeXchange with identifier PXD080179. Source data are provided in Supplementary Table 6.

## Acknowledgements

We thank Ton Timmers and the Central Microscopy facility (CeMic) at MPIPZ for microscopy aid, Aristeidis Stamatakis and the greenhouse team at MPIPZ for help with plant growth, Anne Harzen for MS sample preparation. We are grateful to Stefanie Ranf for providing GoldenGate vectors for molecular cloning and Mark Youles (TSL Norwich) and Laurence Tomlinson (TSL Norwich) for providing the pICSL4723OD vector for CRISPR-Cas9 cloning. We thank Niko Geldner and Cyril Zipfel for providing mutant seeds and for insightful discussions, Michael Hothorn, Cranos Williams and Max Gordon for insightful discussions, and members of the TGA group for their assistance with this project.

## Author contributions

D.S. and T.G.A. conceived the project. D.S., T.G.A., Y.Z., and H.N. designed the experiments. Y.Z. and D.S. performed most of the experiments, with assistance from S.Sa., A.S., S.M., and T.L. J.R. and M.S. created the *Atcep* 12*×* KO mutants. S.St. performed the phosphoproteomic analysis. Y.Z., D.S. and T.G.A. analyzed the data. D.S. and T.G.A. wrote the manuscript with input from all authors. All authors approved the final version of the manuscript.

## Materials & Correspondence

Correspondence and requests for materials should be addressed to Tonni Grube Andersen and Defeng Shen.

## Supplementary Information

Supplementary Table 1. Primers used in this study.

Supplementary Table 2. Differentially expressed genes and enriched GO terms of transcriptomes.

Supplementary Table 3. Differentially phosphorylated peptides identified.

Supplementary Table 4. Canonical nitrogen-responsive genes.

Supplementary Table 5. Acronyms with accession numbers used in this study.

Supplementary Table 6. Source data.

---

**Extended Data Fig. 1.**
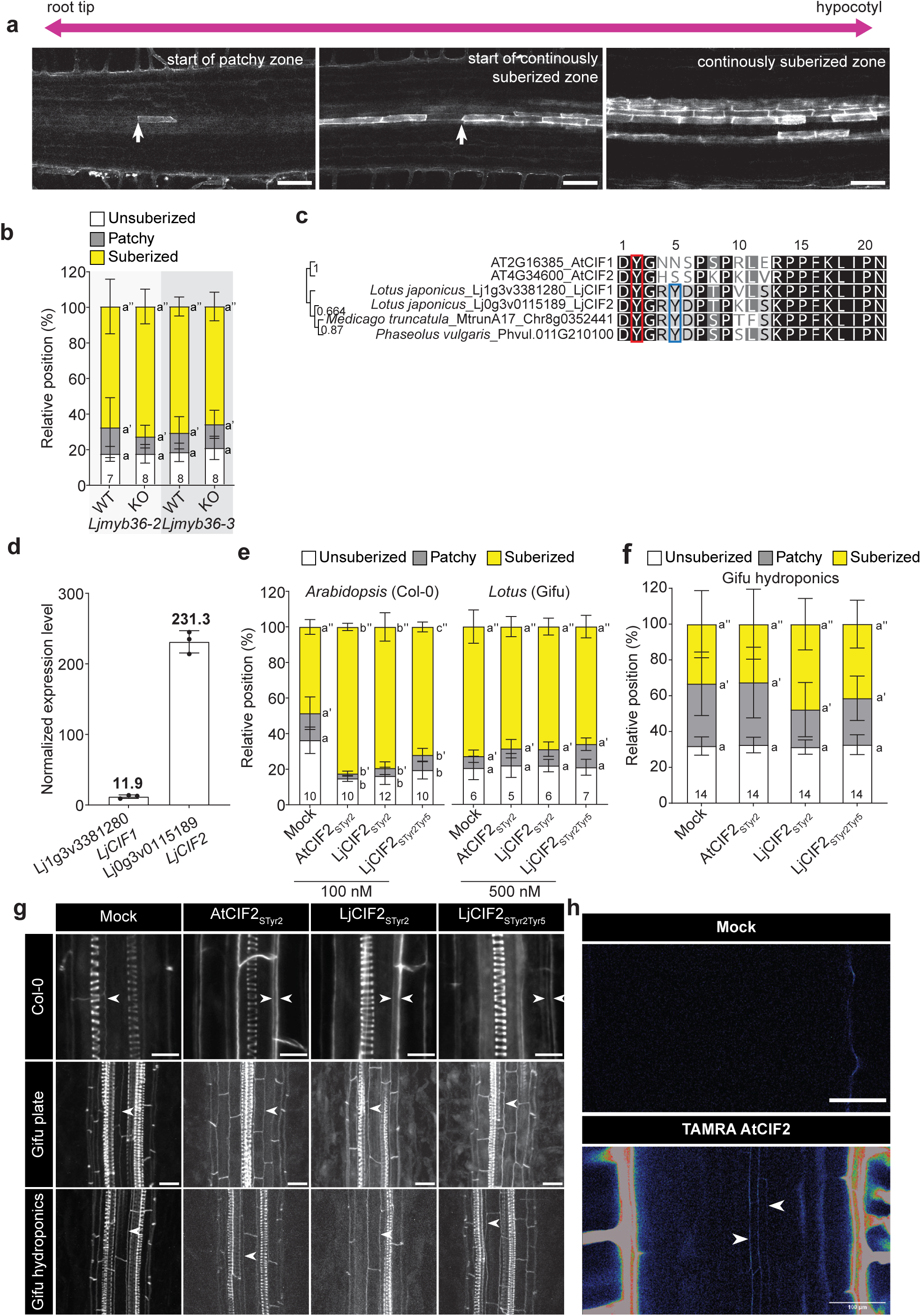
Exogenous CIF peptide does not induce ectopic suberization or lignification in *Lotus* roots. **a**, Representation of suberization pattern in Fluorol Yellow-stained *Lotus* roots. Arrows indicate the start of the patchy suberization zone (left) and the continuously suberized zone (middle). In the continuously suberized zone, only some endodermal cells are continuously suberized. **b**, Suberization patterns in two independent *Ljmyb36* mutant alleles compared with their respective segregating wild type (WT) controls. Different letters depict the statistical difference between mutants with their corresponding segregated WT lines in a two-sided Student’s *t*-test (*P* < 0.05). Representative results from two independent experiments. **c**, Maximum-likelihood phylogenetic tree of CIF peptides from *Arabidopsis*, *Lotus*, *Medicago truncatula* and *Phaseolus vulgaris* based on full-length protein sequences. Nodes containing AtCIF peptides were used as the outgroup. Bootstrap values are indicated at the nodes. Only mature peptide sequences are shown. The red box indicates the conserved Tyr residue at position 2, the blue box highlights a conserved Tyr residue at position 5 that is absent from *Arabidopsis* CIF peptides. **d**, Normalized expression values of *LjCIF1* and *LjCIF2* in *Lotus* roots. Data were obtained from Suzaki *et al.* through *Lotus* Base^49^. Mean expression values are indicated. **e**, Suberization patterns of 7-day-old *Arabidopsis* Col-0 roots (left) treated with 100 nM CIF2 peptide for 48 h under agar conditions and 8-day-old *Lotus* Gifu roots (right) treated with 500 nM CIF2 peptide for 48 h under agar conditions. Representative results from two independent experiments. **f**, Suberization pattern of 8-day-old *Lotus* Gifu roots treated with 500 nM CIF2 peptide for 48 h under hydroponic conditions. **g**, Max projection of confocal image stacks of Basic Fuchsin-stained *Arabidopsis* and *Lotus* roots treated with CIF2 peptides. Double arrowheads indicate overlignification. **h**, Confocal image of an 8-day-old *Lotus* Gifu root treated with 500 nM TAMRA-labelled AtCIF2 peptide for 2 h under hydroponic conditions. Arrowheads indicate penetration of TAMRA-labelled AtCIF2 into the endodermis within the root differentiation zone. Representative image from three roots. Different letters in (**e**) and (**f**) indicate statistically significant differences determined by one-way ANOVA followed by Tukey’s multiple-comparison test (*P* < 0.05). In bar charts, error bars denote standard deviation. The number of biological replicates is indicated in the graphs. Scale bars, 500 µm (**a**), 10 µm for Col-0, 20 µm for Gifu (**g**), and 100 µm (**h**).

**Extended Data Fig. 2.**
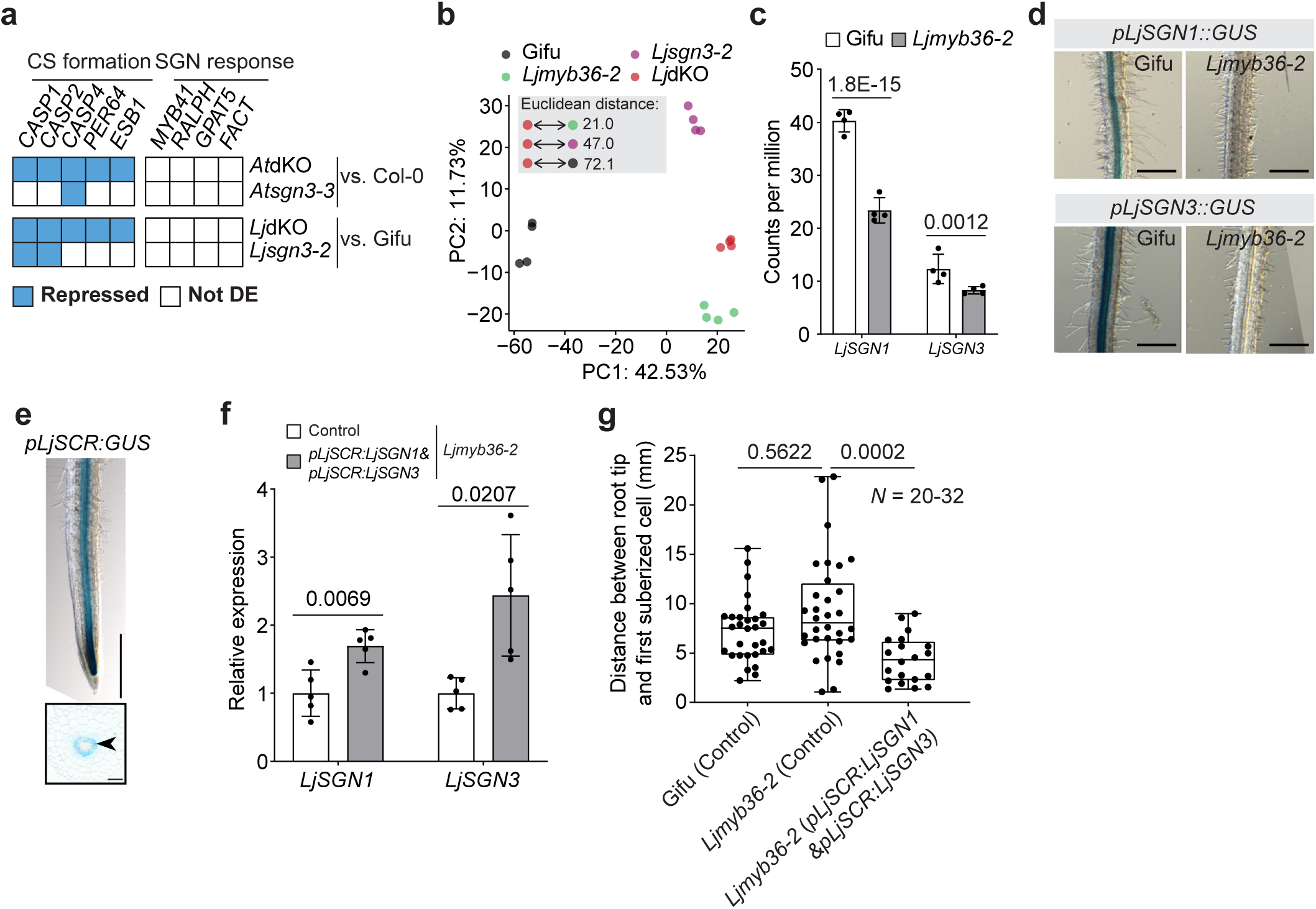
Expression of *SGN1* and *SGN3* in *Lotus* roots requires MYB36. a, Heatmap showing the expression of genes associated with Casparian strip (CS) formation and SGN response in *Arabidopsis* and *Lotus sgn3* and *myb36 sgn3* double knockout (dKO) mutant roots relative to their wild type (WT). Blue: repressed (FDR-adjusted *P* value < 0.05, fold change <-2); white: not differentially expressed (DE). **b**, Principal component analysis of root transcriptome from *Lotus* CS mutants. Euclidean distances between genotypes are shown in the inset. **c**, Normalized counts per million (CPM) of *LjSGN1 and LjSGN3* transcripts in wild type (WT) and *Ljmyb36-2* roots. FDR-adjusted *P* values are indicated. **d**, Transcriptional activity of *LjSGN1* (top) and *LjSGN3* (bottom) in Gifu and *Ljmyb36-2* hairy roots, visualized by GUS signal. Representative images of at least 10 independent composite plants. **e**, *pLjSCR:GUS* activity in Gifu hairy roots. Top, whole-root view; bottom: transverse section through the mature root zone. Representative images from 10 independent composite plants. Arrowhead indicates the endodermis. **f**, Relative expression levels of *LjSGN1* and *LjSGN3* in *Ljmyb36-2* hairy roots expressing *pLjSCR:GUS* (Control) or *pLjSCR:LjSGN1&pLjSCR:LjSGN3*. Statistical difference was determined by two-sided Student’s *t*-test. **g**, Distance between the root tip and the first suberized endodermal cell in *Ljmyb36-2* hairy roots overexpressing *LjSGN1* and *LjSGN3* using *pLjSCR*, compared with transgenic controls (EV, *pLjSCR:GUS*) of Gifu and *Ljmyb36-2*. Combined results of two independent experiments. Statistical difference was determined by two-sided Student’s *t*-test. In bar charts, error bars denote standard deviation. In box plots, the center line indicates the median, box limits represent the 25th and 75th percentiles, and whiskers extend to the most extreme data points. Individual values are shown as scattered jittered dots. The number of biological replicates is indicated on the graph. Scale bar: 500 µm (**d**, **e**, top), 50 µm (**e**, bottom).

**Extended Data Fig. 3.**
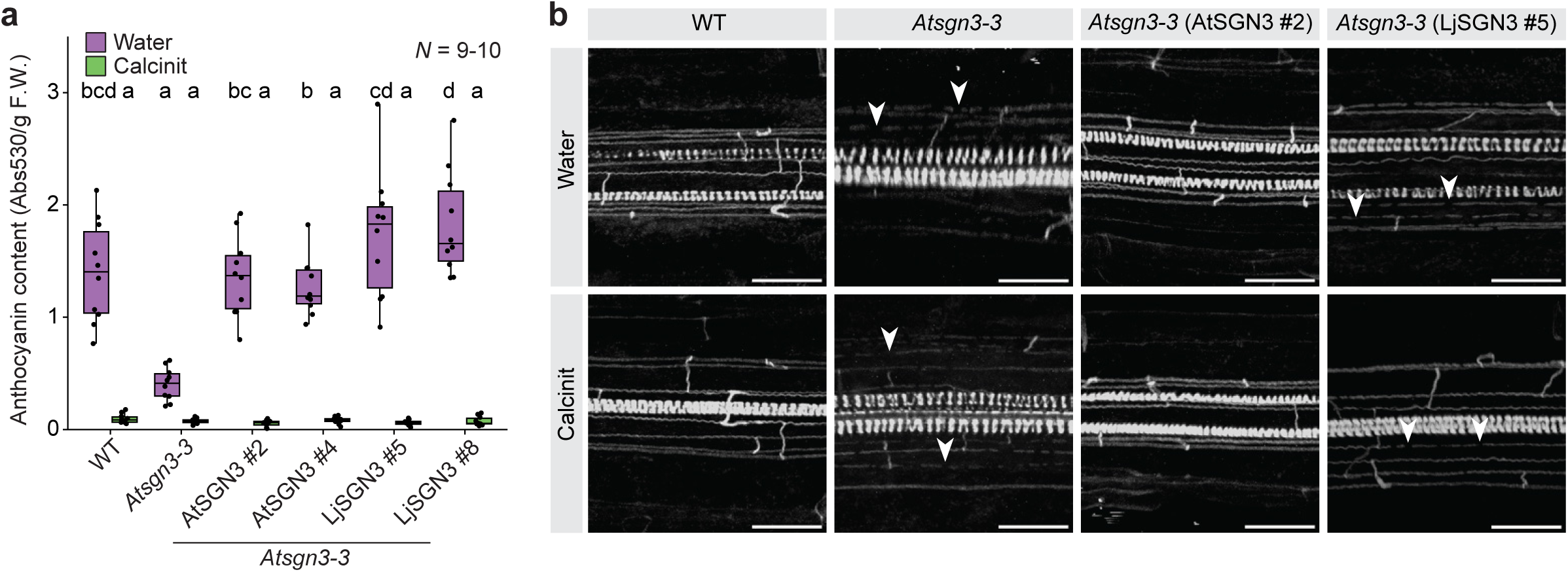
LjSGN3 restores shoot nitrogen response but not root barrier function. **a**, Anthocyanin content in shoots of plants grown in Cologne agricultural soil (CAS) under water-treated or nitrogen-supplemented (Calcinit) conditions. Different letters indicate statistically significant differences determined by one-way ANOVA followed by Sidak’s multiple-comparison test (*P* < 0.05). **b,** Basic Fuchsin-stained roots of plants grown in CAS under water-treated or Calcinit conditions. Arrowheads indicate discontinuities in the Casparian strip. In box plots, the center line indicates the median, box limits represent the 25th and 75th percentiles (IQR), and whiskers extend to the most extreme data points within 1.5 × IQR. Individual values are shown as scattered jittered dots. The number of biological replicates is indicated in the graphs. Scale bar, 25 μm.

**Extended Data Fig. 4.**
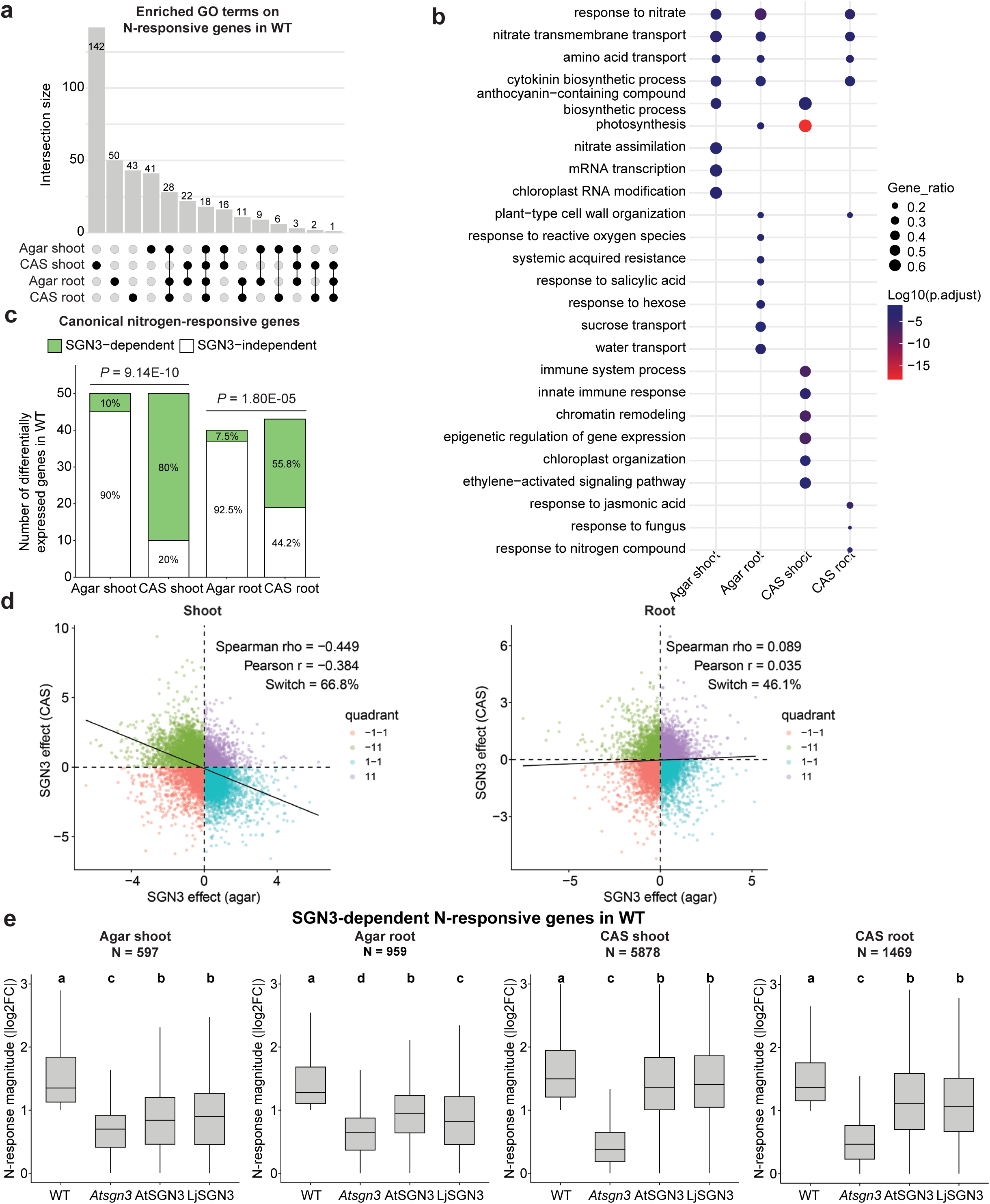
SGN3-dependent nitrogen signaling is conserved across species but undergoes condition- and tissue-specific transcriptional rewiring. **a**, UpSet plot showing limited overlap among nitrogen-responsive genes identified in wild type (WT) shoots and roots grown on agar or in Cologne agricultural soil (CAS). **b**, Dot plot showing distinct Gene Ontology terms enriched among nitrogen-responsive genes in WT shoots and roots under agar and CAS conditions. **c**, Bar plot showing that a significantly greater proportion of canonical nitrogen-responsive genes are SGN3-dependent in WT shoots and roots under CAS conditions than under agar conditions (Full lists in Supplementary Table 4). Statistical significance was determined using Fisher’s exact test. **d**, Scatter plots showing shoot (left) and root (right) SGN3-dependent transcriptional effects (WT – *sgn3* logFC difference) in agar (x-axis) and CAS (y-axis) conditions for each gene. Each point represents a gene, colored by quadrant according to the direction of response in the two environments. Dashed lines indicate zero effect in each condition. The black line shows the least-squares linear regression fit. Correlation structure was assessed using both Spearman and Pearson tests, and the fraction of genes switching response direction between conditions is indicated. **e**, Both AtSGN3 and LjSGN3 restore SGN3-dependent nitrogen responses in WT shoots and roots under agar and soil conditions. Different letters indicate statistically significant differences determined by one-way ANOVA followed by Tukey’s multiple-comparison test (*P* < 0.05). Numbers shown in the graph indicate the number of SGN3-dependent nitrogen-responsive genes identified in WT plants. In box plots, the center line indicates the median, box limits represent the 25th and 75th percentiles (IQR), and whiskers extend to the most extreme data points within 1.5 × IQR.

**Extended Data Fig. 5.**
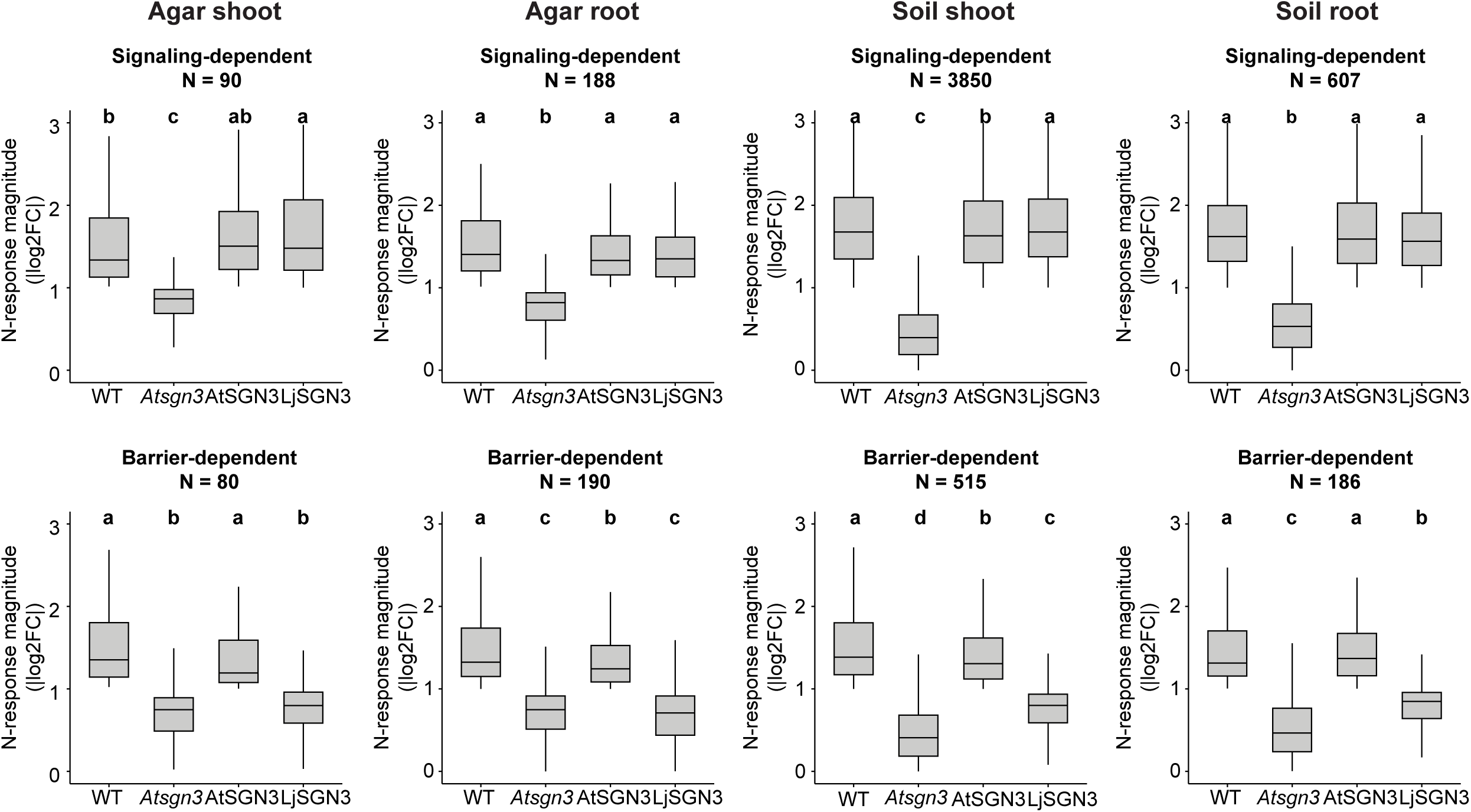
Categorization of signaling- and barrier-dependent nitrogen-responsive genes in wild type shoots and roots under agar and soil conditions. Nitrogen-responsive genes in wild type (WT) plants were first identified based on differential expression upon nitrogen treatment. These genes were then classified into two subcategories according to their expression patterns across genotypes: signaling-dependent nitrogen-responsive genes were defined as those that lost nitrogen responsiveness in the *Atsgn3-3* mutant but regained it upon complementation with either *AtSGN3* or *LjSGN3*; barrier-dependent nitrogen-responsive genes were defined as those that lost responsiveness in *Atsgn3-3* and were restored by *AtSGN3* but not by *LjSGN3*. Different letters indicate statistically significant differences determined by one-way ANOVA followed by Tukey’s multiple-comparison test (*P* < 0.05). Numbers shown in the graph indicate the number of nitrogen-responsive genes belonging to each category. In box plots, the center line indicates the median, box limits represent the 25th and 75th percentiles (IQR), and whiskers extend to the most extreme data points within 1.5 × IQR.

**Extended Data Fig. 6.**
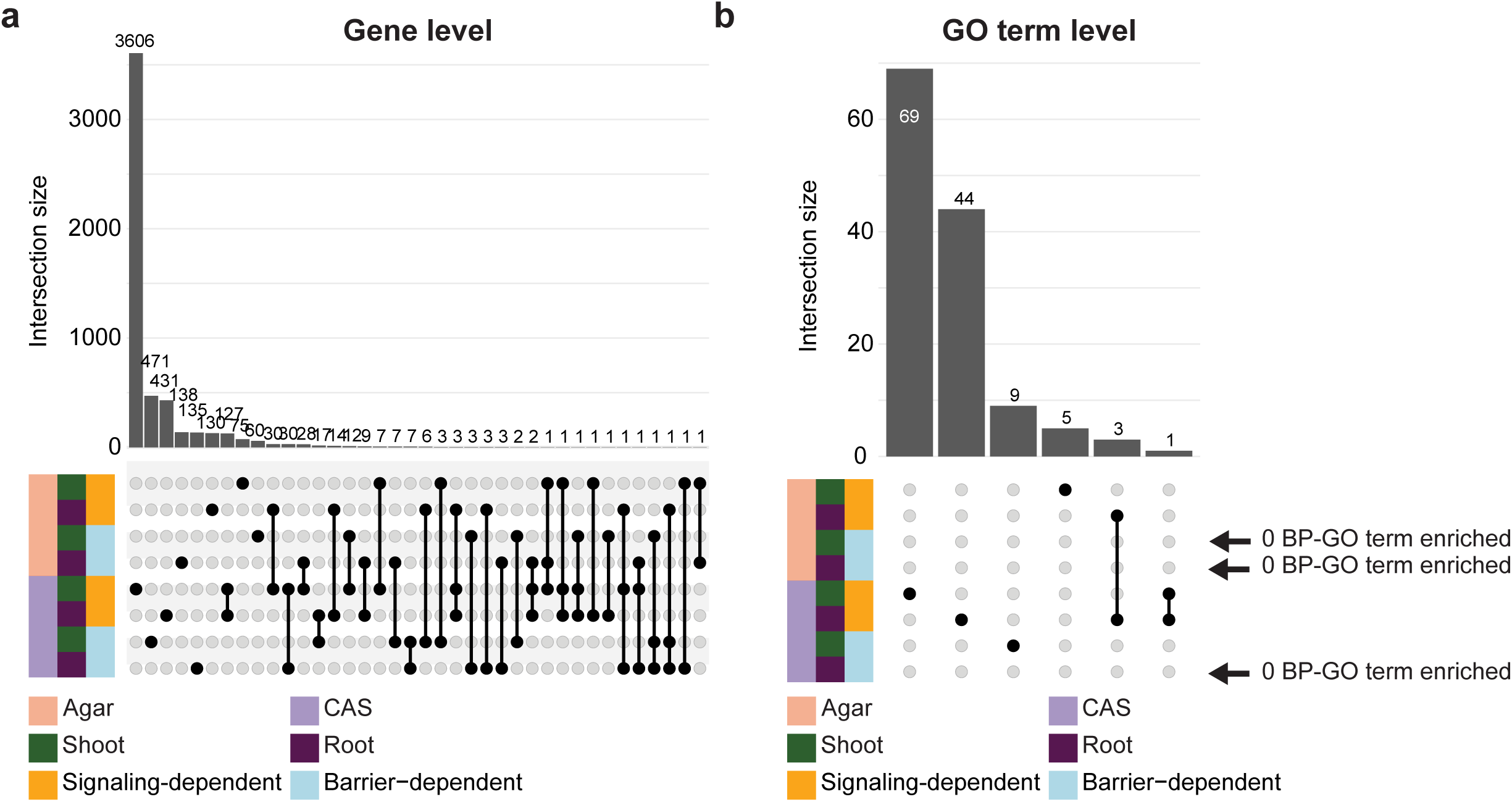
Limited overlap among signaling-dependent and barrier-dependent nitrogen responses across tissues and growth conditions. a, UpSet plot showing the overlap among signaling-dependent and barrier-dependent nitrogen-responsive genes identified in wild type (WT) shoots and roots grown on agar or in Cologne agricultural soil (CAS). **b,** UpSet plot showing the overlap among Biological Process (BP) Gene Ontology (GO) terms enriched in signaling-dependent and barrier-dependent nitrogen responses across tissues and growth conditions.

**Extended Data Fig. 7.**
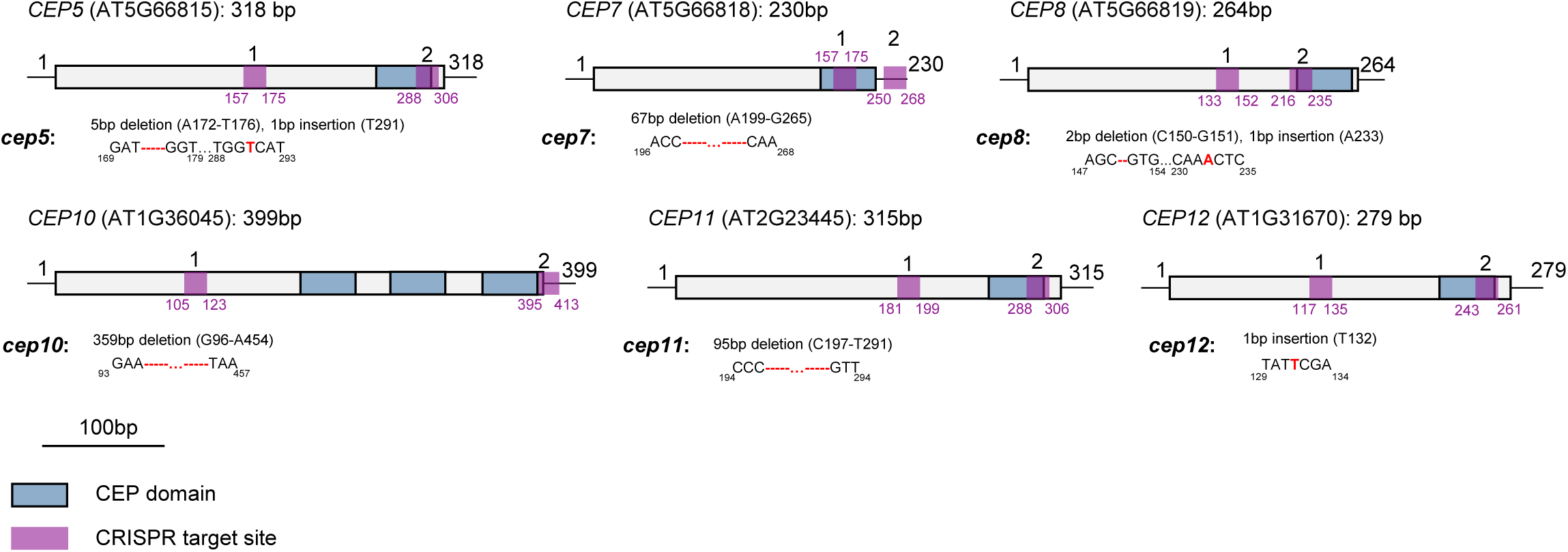
Generation and mutagenesis information for the *Atcep* 12× knockout mutant. The *Atcep* 12× knockout (KO) line was generated by introducing additional targeted mutations into the previously established *Atcep* 6× KO background^26^. Schematic diagram of *CEP5*, *CEP7*, *CEP8*, *CEP10*, *CEP11* and *CEP12* gene structure and the CRISPR-Cas9-mediated mutation pattern detected by DNA sequencing. The locus number and the length of the coding sequence (CDS) are indicated above the scheme for each gene. The specific location and type of mutations for each gene are indicated in the schematics describing the mutants. The CEP domain is indicated in blue, and the two CRISPR target sites are indicated in purple. Scale bar, 100 bp.

**Extended Data Fig. 8.**
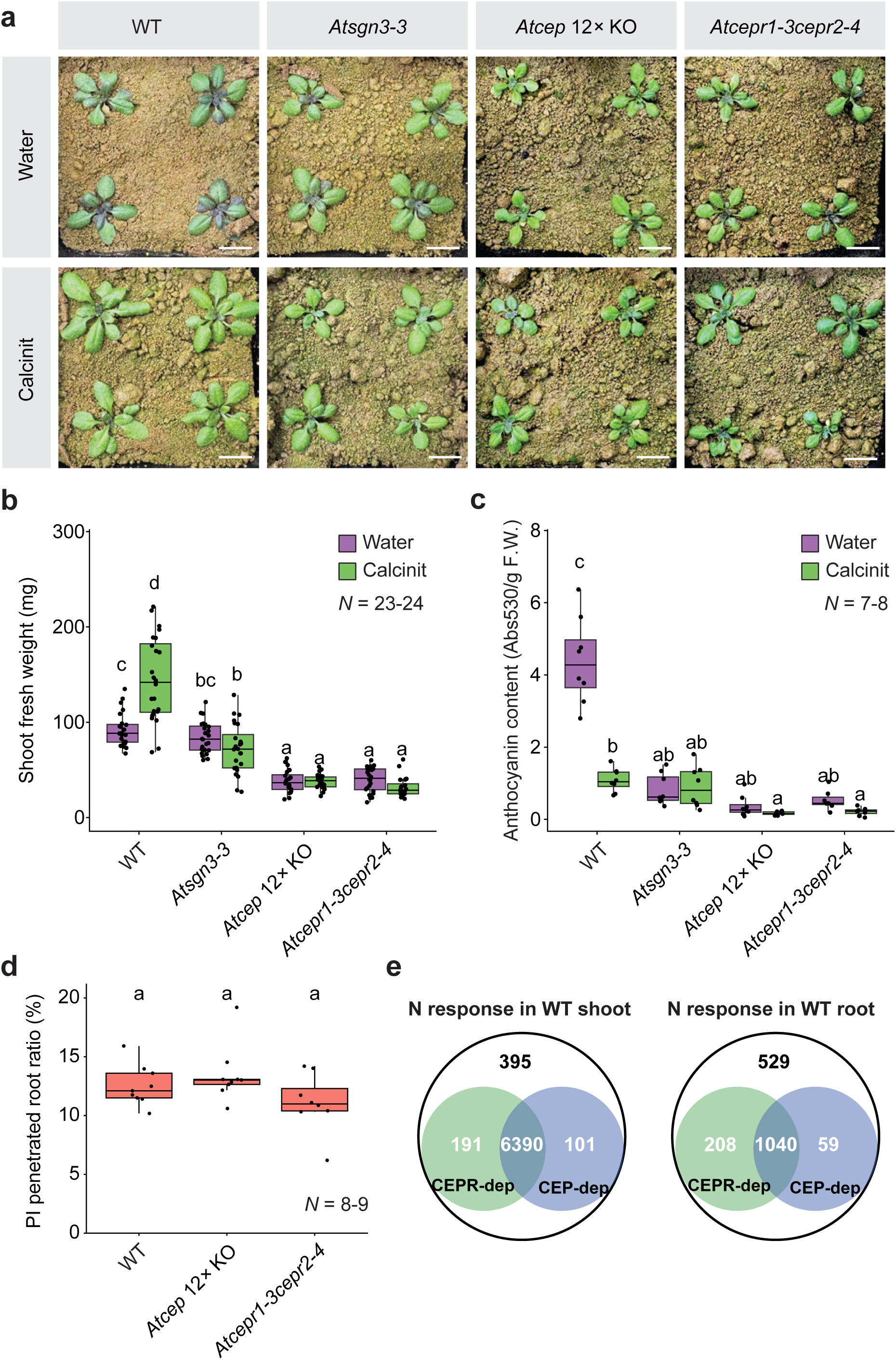
Nitrogen responses in wild type under soil conditions are largely CEP–CEPR dependent. a, Representative images of 21-day-old plants grown in Cologne agriculture soil (CAS) under water-treated and nitrogen-supplemented (Calcinit) conditions. **b,c**, Shoot fresh weight (**b**) and anthocyanin content (**c**) in shoots of plants grown in CAS under water-treated or Calcinit conditions. **d**, Proportion of roots exhibiting propidium iodide (PI) penetration in 7-day-old seedlings. **e**, Venn diagram depicting the number of nitrogen-responsive genes in WT that are CEPR-dependent and CEP-dependent. Different letters in (**b, c**) indicate statistically significant differences determined by one-way ANOVA followed by Sidak’s multiple-comparison test (*P* < 0.05). Different letters in (**d**) indicate statistically significant differences determined by one-way ANOVA followed by Tukey’s multiple-comparison test (*P* < 0.05). In box plots, the center line indicates the median, box limits represent the 25th and 75th percentiles (IQR), and whiskers extend to the most extreme data points within 1.5 × IQR. Individual values are shown as scattered jittered dots. The number of biological replicates is indicated on the graph. Scale bar, 1 cm.

**Extended Data Fig. 9.**
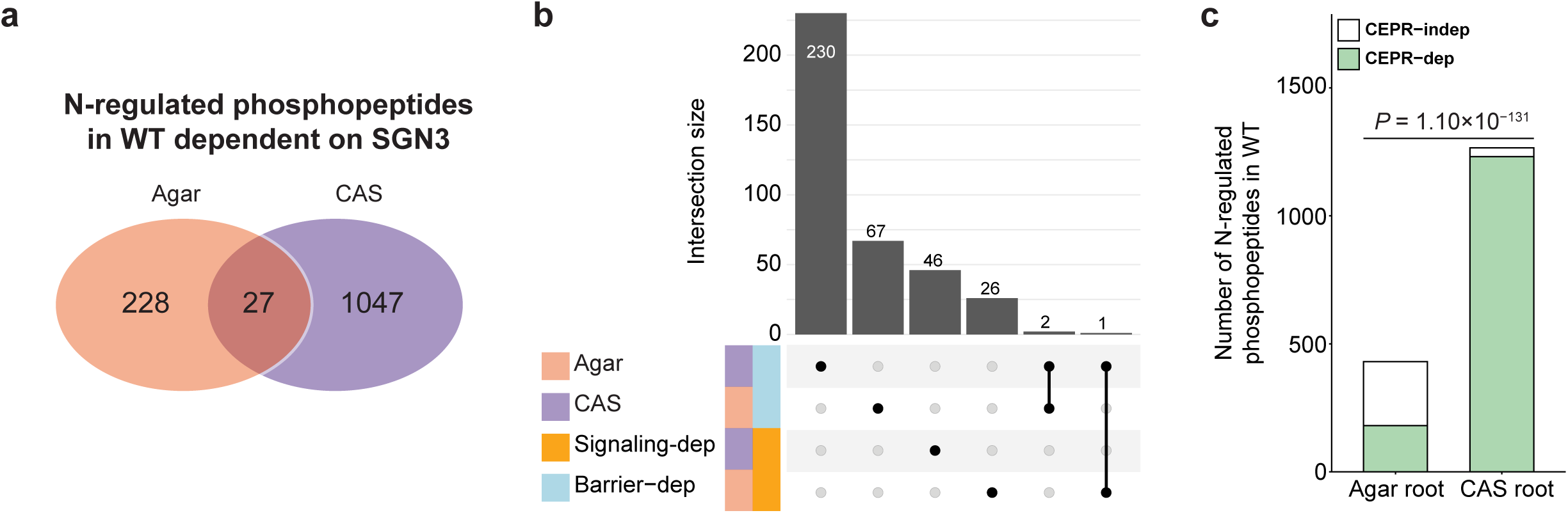
Phosphoproteomic responses in wild type roots differ between agar and soil conditions. **a**, Venn diagram showing limited overlap between SGN3-dependent phosphopeptide changes detected under agar and Cologne agricultural soil (CAS) conditions. **b**, Upset plot showing limited overlap among Gene Ontology (GO) terms enriched in signaling-dependent and barrier-dependent phosphosite changes. **c**, Distribution of nitrogen-regulated phosphopeptides across different categories in wild type (WT) roots grown under agar or CAS conditions. Fisher’s exact test was used to compare the proportion of CEPR-dependent phosphopeptide changes between agar and CAS conditions.

**Extended Data Fig. 10.**
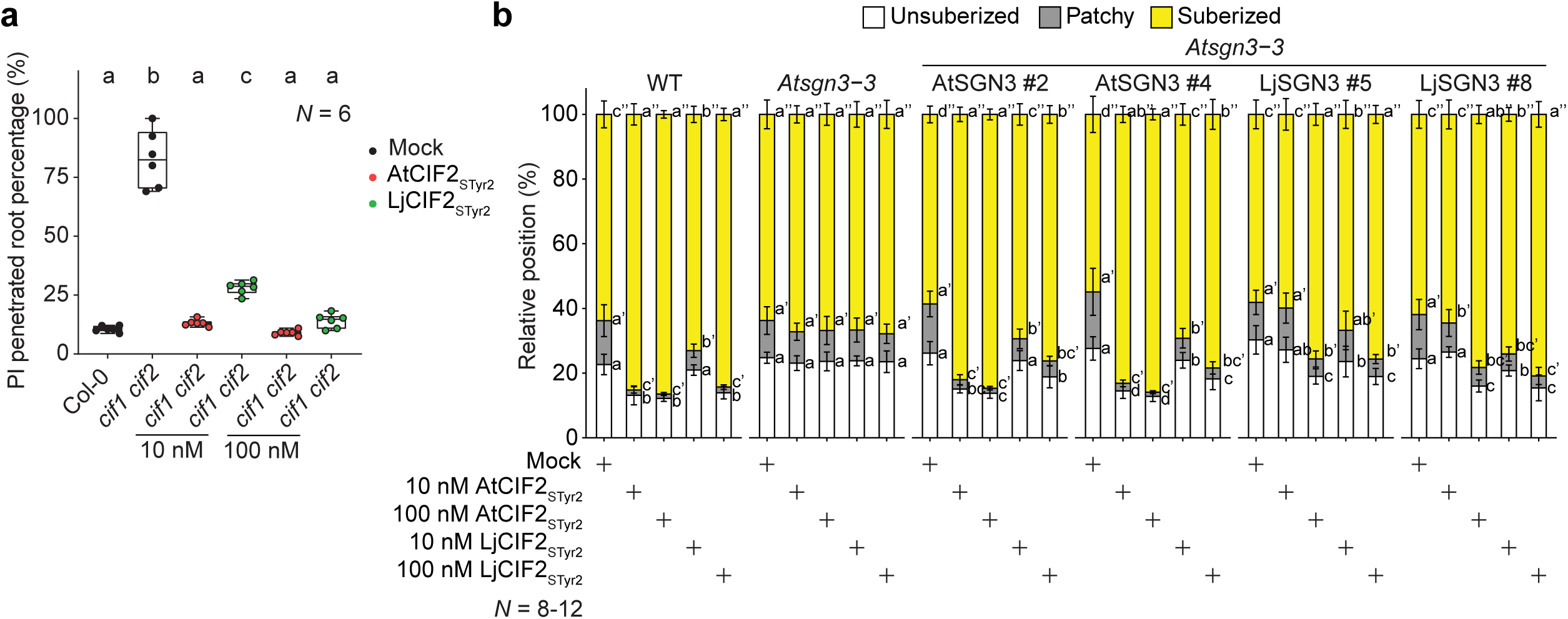
SGN3 is more responsive to native ligands in inducing Casparian strip formation and ectopic suberization. **a**, Proportion of roots exhibiting propidium iodide (PI) penetration in 7-day-old seedlings after 2 days of CIF2 peptide treatment. Different letters indicate statistically significant differences determined by one-way ANOVA followed by Tukey’s multiple-comparison test (*P* < 0.05). **b**, Suberization patterns of 7-day-old *Atsgn3-3* complementation lines in response to AtCIF2 and LjCIF2 peptides after 2 days of CIF2 peptide treatment. Different letters indicate statistically significant differences determined by one-way ANOVA followed by Tukey’s multiple-comparison test (*P* < 0.05) within the same genotype. In bar charts, error bars denote standard deviation. In box plots, the center line indicates the median, box limits represent the 25th and 75th percentiles (IQR), and whiskers extend to the most extreme data points within 1.5 × IQR. Individual values are shown as scattered jittered dots. The number of biological replicates is indicated in the graphs.

**Extended Data Fig. 11.**
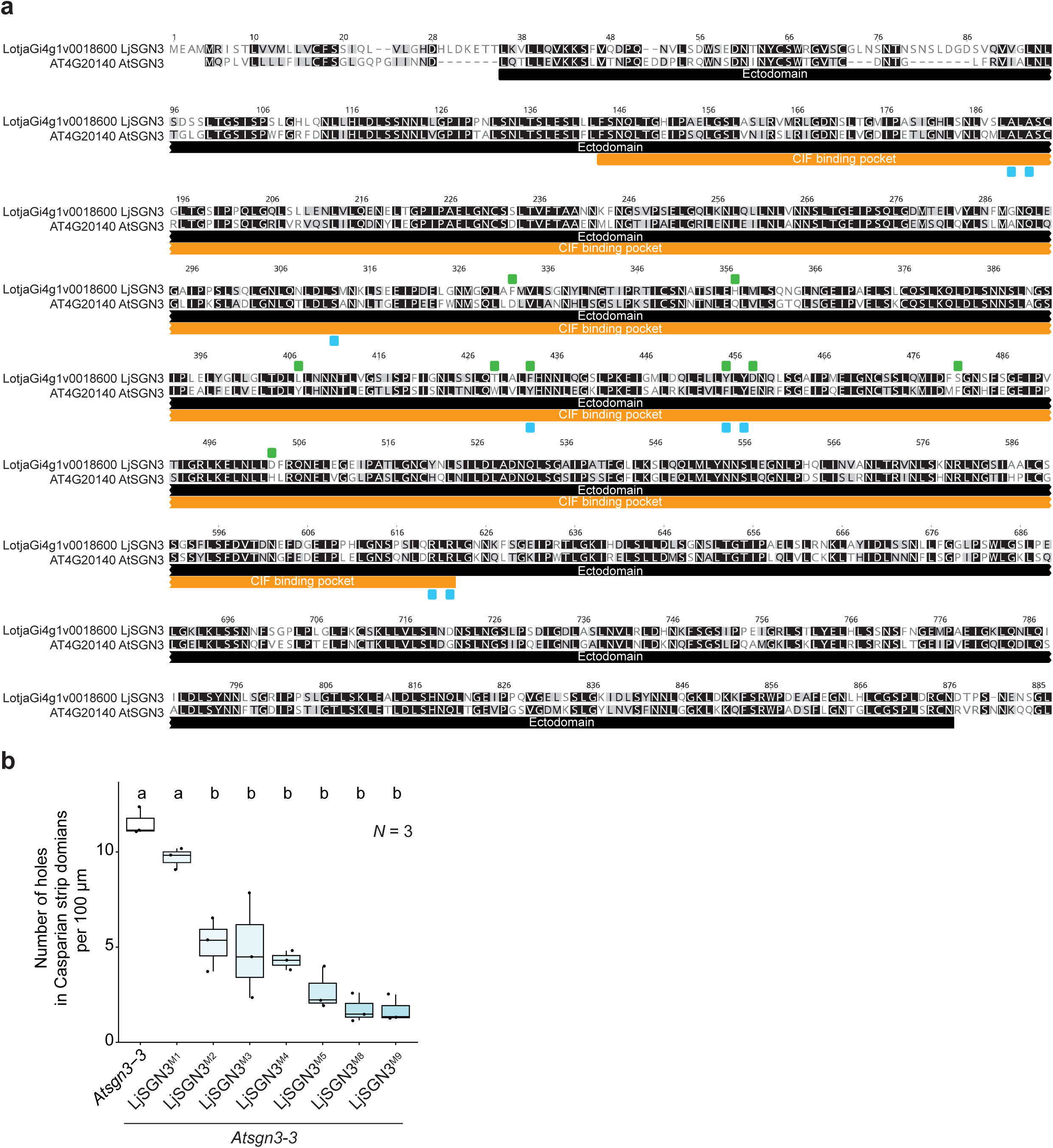
Functional complementation of the LjSGN3 ectodomain by AtSGN3 residues rescues Casparian strip continuity. a, Protein sequence alignment of the N-terminal region of LjSGN3 and AtSGN3. The ectodomain is referenced to the crystal structure of AtSGN3 ectodomain, with structurally conserved regions indicated by black boxes. The CIF-binding pocket of AtSGN3 is highlighted by orange boxes. Previously experimentally characterized residues in AtSGN3 are marked with blue boxes. while the LjSGN3 residues that were mutagenized to their AtSGN3 counterparts are denoted with green boxes. Amino acid coordinates are numbered according to the LjSGN3 sequence. **b**, Quantification of Casparian strip discontinuities, measured as the number of holes in the Casparian strip domain per 100 µm of endodermis in 9-day-old seedlings. Different letters indicate statistically significant differences determined by one-way ANOVA followed by Tukey’s multiple-comparison test (*P* < 0.05). In box plots, the center line indicates the median, box limits represent the 25th and 75th percentiles (IQR), and whiskers extend to the most extreme data points within 1.5 × IQR. Individual values are shown as scattered jittered dots. The number of biological replicates is indicated in the graphs.

**Extended Data Fig. 12.**
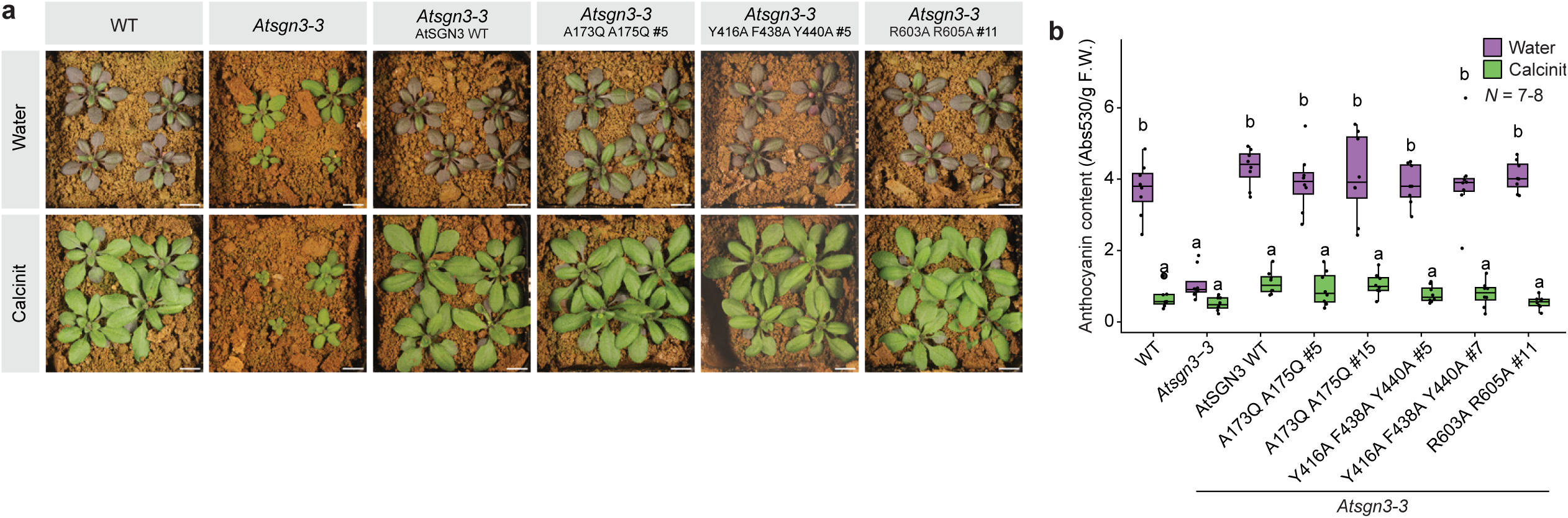
SGN3 variants with lower CIF2 binding affinity restore shoot nitrogen response. **a**, Representative images of 28-day-old plants grown in CAS under water-treated and Calcinit conditions. The number of biological replicates is indicated in the graphs. **b**, Anthocyanin content in shoots of plants grown in Cologne agriculture soil (CAS) under water-treated or nitrogen-supplemented (Calcinit) conditions. Different letters indicate statistically significant differences determined by one-way ANOVA followed by Sidak’s multiple-comparison test (*P* < 0.05). In box plots, the center line indicates the median, box limits represent the 25th and 75th percentiles (IQR), and whiskers extend to the most extreme data points within 1.5 × IQR. Individual values are shown as scattered jittered dots. The number of biological replicates is indicated in the graphs. Scale bar, 1 cm.

**Extended Data Fig. 13.**
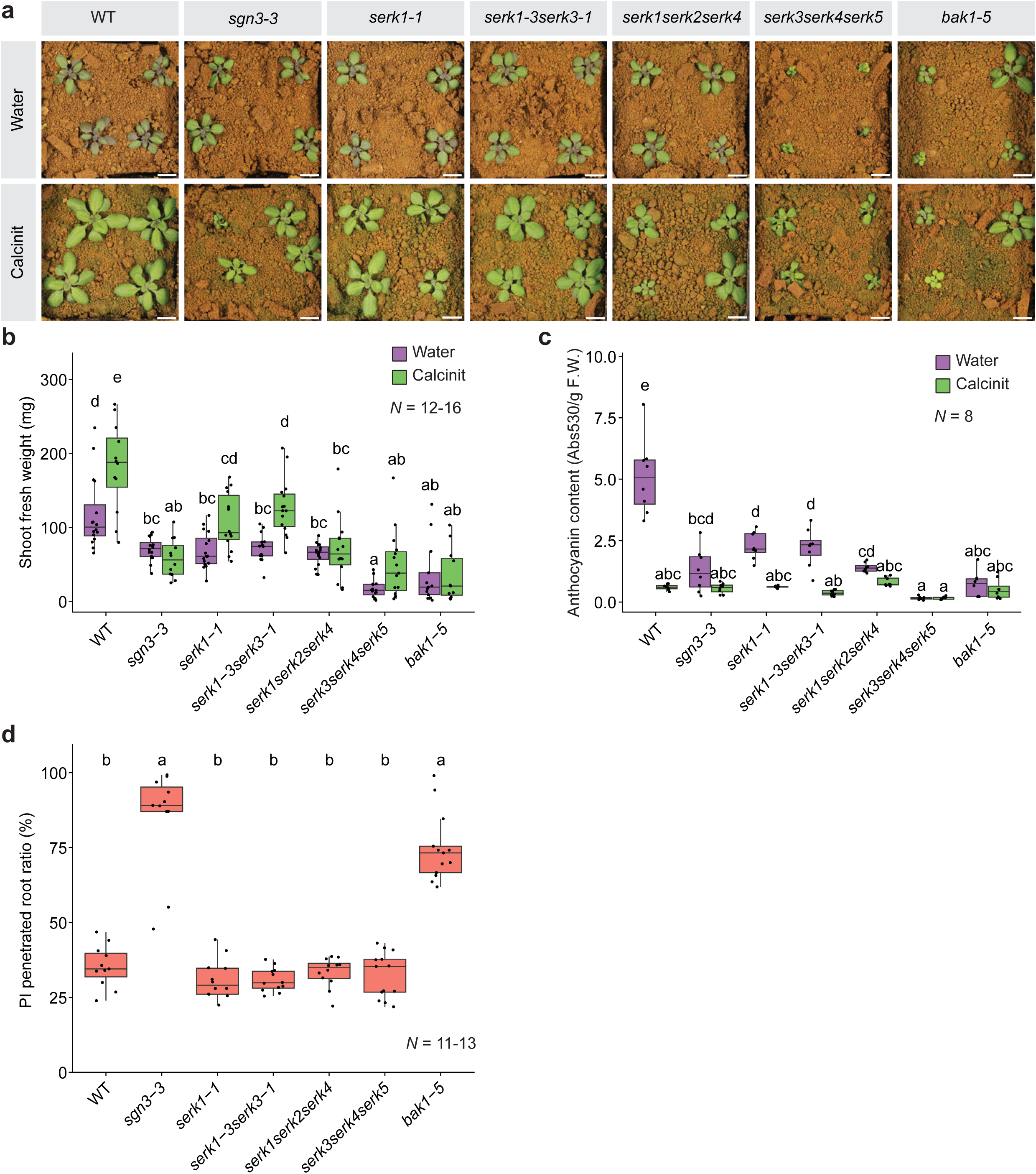
*serk1* and high-order *serk* mutants display diminished shoot nitrogen response while maintaining intact Casparian strip function. **a**, Representative images of 21-day-old plants grown in Cologne agriculture soil (CAS) under water-treated and nitrogen-supplemented (Calcinit) conditions. **b,c**, Shoot fresh weight (**b**) and shoot anthocyanin content (**c**) plants grown in CAS under water-treated or Calcinit conditions. **d**, Proportion of roots exhibiting propidium iodide (PI) penetration in 7-day-old seedlings. Different letters in (**b,c**) indicate statistically significant differences determined by one-way ANOVA followed by Sidak’s multiple-comparison test (*P* < 0.05). Different letters in (**d**) indicate statistically significant differences determined by one-way ANOVA followed by Tukey’s multiple-comparison test (*P* < 0.05). In box plots, the center line indicates the median, box limits represent the 25th and 75th percentiles (IQR), and whiskers extend to the most extreme data points within 1.5 × IQR. Individual values are shown as scattered jittered dots. The number of biological replicates is indicated in the graphs. Scale bar, 1 cm.

**Extended Data Fig. 14.**
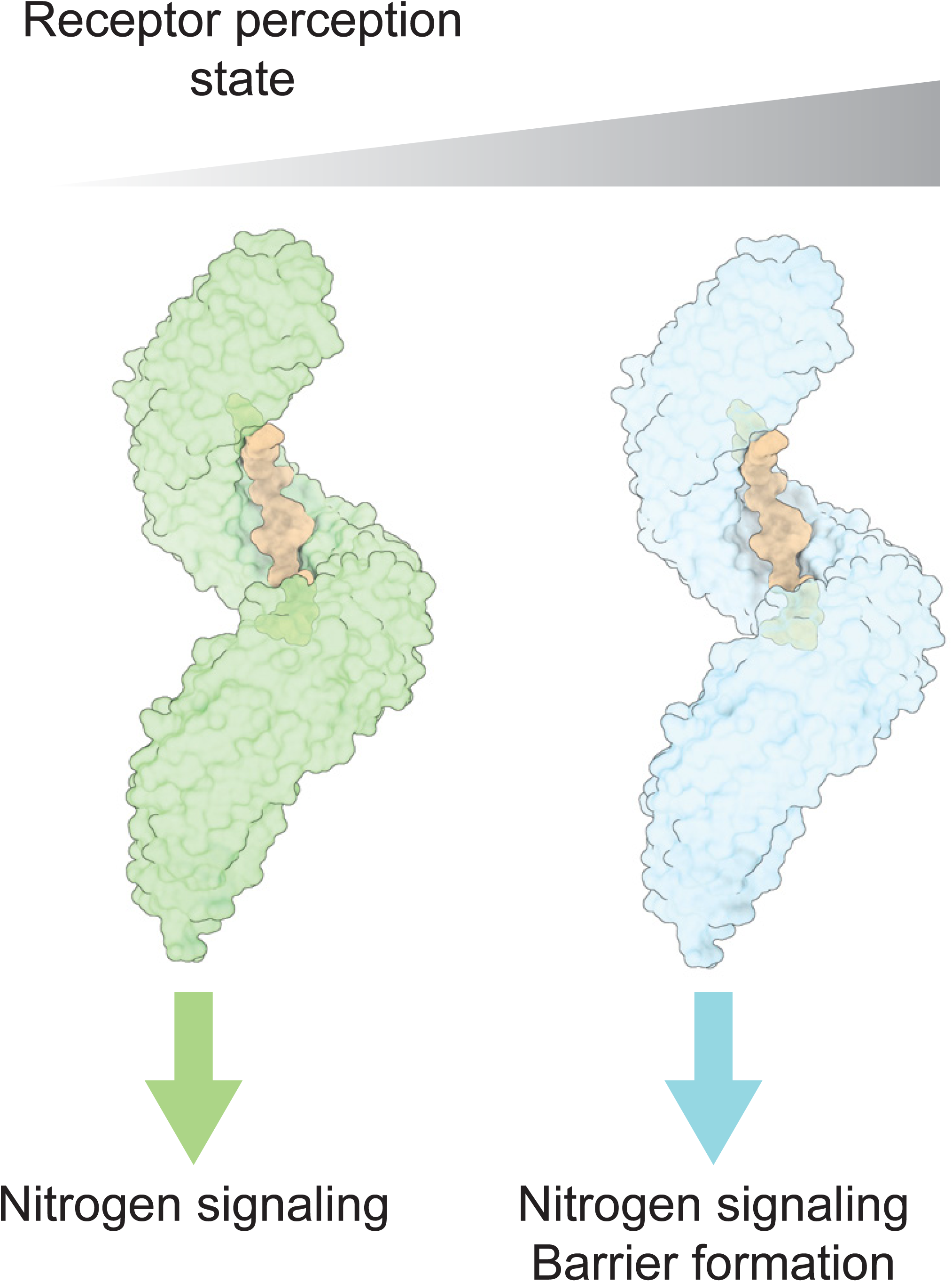
Proposed model illustrating how SGN3 perception state biases distinct outputs. A low receptor perception state preferentially promotes nitrogen signaling, whereas a high receptor perception state directs both nitrogen signaling and barrier formation. The ligand is shown in orange. Receptors in low and high perception states are shown in light green and light blue, respectively.

